# Three-dimensional actuation of hummingbird wings requires diverse effects from the primary flight muscles

**DOI:** 10.1101/2022.01.27.477984

**Authors:** Suyash Agrawal, Bret W. Tobalske, Zafar Anwar, Haoxiang Luo, Tyson L. Hedrick, Bo Cheng

## Abstract

Hummingbirds have evolved to hover and maneuver with exceptional flight control. This is directly enabled by their musculoskeletal system that successfully exploits the agile motion of flapping wings. Here, we reveal novel principles of hummingbird wing actuation that provide insights into the evolution and robotic emulation of hummingbird flight. We develop a functional model of hummingbird musculoskeletal system, which predicts instantaneous, three-dimensional torque produced by primary (pectoralis and supracoracoideus) and combined secondary muscles. It also reveals primary muscle contractile behavior, including stress, strain, elasticity, and work. Results show that the primary muscles (i.e., the flight “engine”) function as diverse effectors, as they do not simply power the stroke, but also actively deviate and pitch the wing. The secondary muscles produce controlled-tightening effects, by acting against primary muscles in deviation and pitching. The diverse effector capacity of pectoralis is associated with the evolution of a comparatively enormous bicipital crest on humerus.

## Introduction

Hummingbirds use their flapping wings to control aerodynamic forces and moments to an extent unmatched by other biological and robotic fliers [1–7]. Such high authority of flight control is directly enabled by their wing musculoskeletal system [8] (Fig. 1). The hummingbird wing skeleton has anatomical components [9,10] and degrees of freedom similar to those of other flying birds (Fig. 1a); however, the proportions of different components have evolved convergently to that of insect wings to create an approximately three-degree-of-freedom (3-DoF) wing motion about the shoulder joint [8,11] (stroke, deviation, and pitching, Fig. 1b), with gearing of the primary power muscles (pectoralis and supracoracoideus) similar to insects. However, the anatomical and physiological features of hummingbird wings do not simply translate into an expressive biomechanical model that can directly reveal the key actuation principles of hummingbird wings, especially as the small size of hummingbirds prevents most *in-vivo* measurements of muscle contractile behavior. As a result, even though the charismatic flight capacity of hummingbirds has attracted much research, a comprehensive understanding of the functioning of hummingbird wing musculoskeletal system is lacking.

**Fig. 1.**
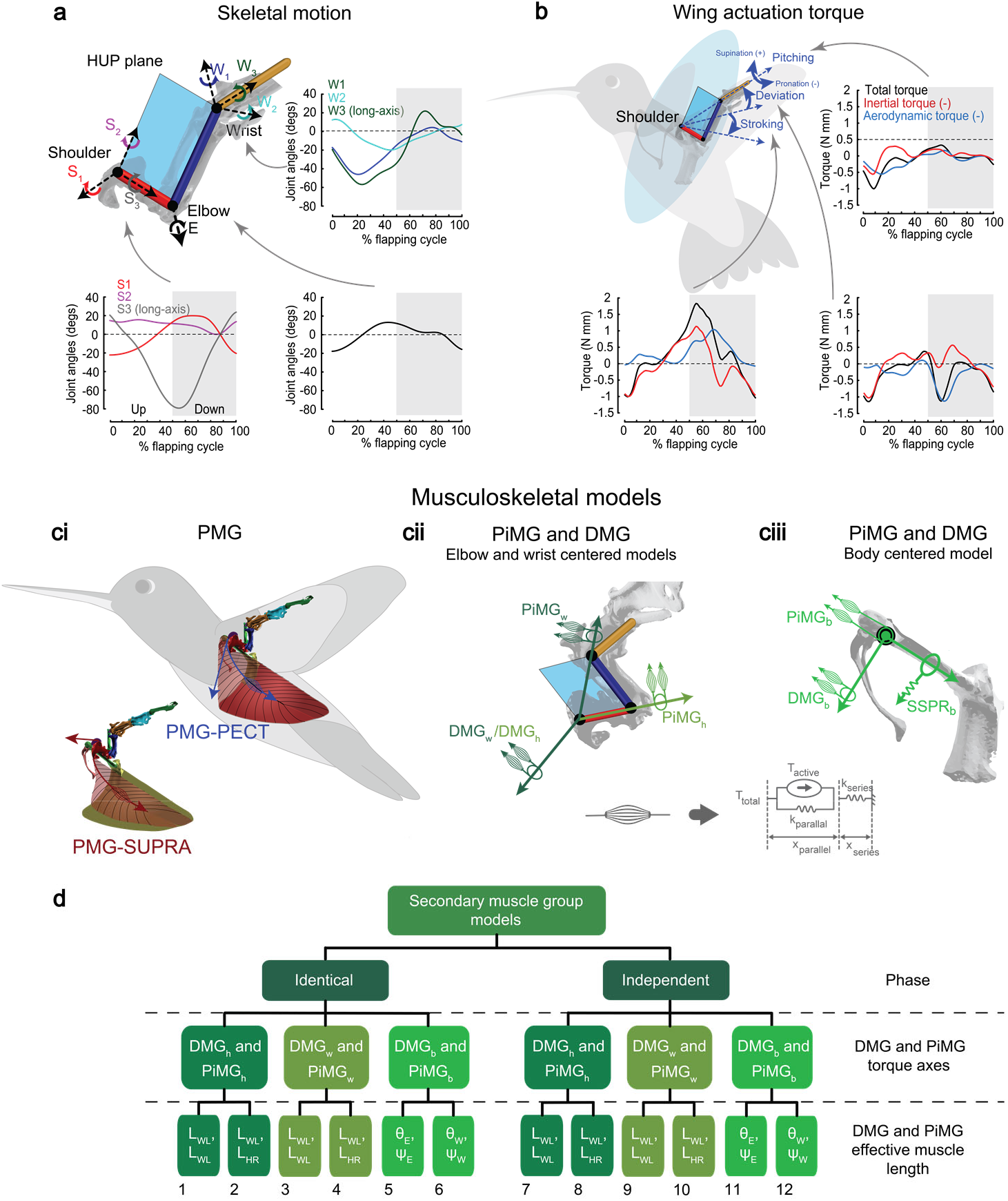
Hummingbird wing skeletal motion, muscle actuation torque, and hypothesized wing-actuation model and its variations. **(a)** Hummingbird wing skeleton as a 7-DoF, functional 3-link system and the definition of respective joint angles. The three functional links include humerus (red), radius and ulna (blue), and carpus, metacarpus, and digits (yellow). The shoulder and wrist act as a 3-DoF ball-and-socket joints and the elbow acts as a 1-DoF hinge joint. The graphs show the instantaneous joint angles over a flapping cycle for the shoulder, elbow, and wrist. The angles were calculated in reference to the wing orientation at the mid-downstroke [8]. Gray shaded and unshaded areas represent downstroke and upstroke, respectively. **(b)** Wing stroke, deviation, and pitching angles defined according to the movement of Humerus-Ulna Plane (HUP, see (a)). The graphs show the corresponding wing muscle actuation torque via the shoulder for each angle (estimated based on the CFD data [19]). Together the wing muscle actuation torque (b) and the wing kinematics (a) were used to optimize the parameters of the hypothesized wing-actuation model (c). **(c)** Primary and secondary muscle groups defined in the hypothesized wing-actuation model. **(ci)** Primary Muscle Group (PMG), including Pectoralis: PMG-PECT and Supracoracoideus: PMG-SUPRA. The muscles were assumed to lie effectively along the aponeurosis arcs and the forces were assumed to act tangent to these arcs at the humerus insertion points. The blue and red straight arrows represent the effective muscle force direction at a representative time instant. Hypothesized actuation axes for secondary muscle groups that include Deviation Muscle Group (DMG) and Pitching Muscle Group (PiMG) for **(cii)** elbow-centered and wrist-centered model variations, and **(ciii)** body-centered model variations. Note that while the muscles have been shown to be antagonistic, we provided the provision for the active torques due to the two muscles in DMG (or PiMG) to act along identical or opposite directions. Also shown below is the three-component model of an individual muscle. **(d)** Summary of the 12 model variations with three layers of assumptions on the right. Model numbers are shown at the bottom. See ‘Methods’ for the muscle effective length definitions.

For engineers, while it is enticing to emulate hummingbirds for the design of novel micro aerial vehicles with high maneuverability and stability, the lack of understanding in hummingbird wing actuation presently renders such bio-mimicry elusive and unfruitful. Existing hummingbird-inspired or hummingbird-sized flapping-wing robot designs (with hovering capacity) [7,12–18] replicate only a few hypothetical hummingbird flight features, such as stroking using single-axis wing actuation, high elastic energy storage, and passive wing pitching. In addition, the existing designs do not explicitly consider other flight modes that undoubtedly have also shaped the evolution of the biomechanical design of hummingbird wings. Therefore, it is unlikely that these engineering designs have captured the key phenotypic traits that are needed to emulate the complete envelope and capacity of hummingbird flight ranging from hovering [8] to various agile maneuvers that do not conform to the helicopter models [1].

We provide novel insights into the biomechanics of hummingbird wing actuation via functional modeling grounded in the complexities of musculoskeletal and kinematic processes. The hummingbird wing musculoskeletal system effectively comprises of three rigid links connected in series (Fig. 1a), giving rise to a 7-DoF skeletal system. Based on the available data on hummingbird muscle anatomy [9,10] we found that the muscles connecting the body skeleton to the wing skeleton primarily act along the rigid Humerus-Ulna Plane (HUP, Fig. 1a). Our hypothesized functional models for the hummingbird wing musculoskeletal system (referred to below as musculoskeletal wing-actuation models), therefore, were comprised of three antagonistic muscle groups that provided actuation to HUP about the shoulder joint, generating an approximate 3DoF wing motion (Fig. 1b and c). These include a Primary Muscle Group (PMG) consisting of two primary muscles (Fig. 1ci), pectoralis and supracoracoideus (referred to as PECT and SUPRA in the wing-actuation model), and two combined secondary muscle groups, including Deviation Muscle Group (DMG) and Pitching Muscle Group (PiMG)) (Fig. 1cii and ciii). We then developed a total of twelve variations of the wing-actuation model with identical insertion points of primary muscles, but different assumptions on the secondary muscle groups (Fig. 1c and d). The functional model for each muscle includes an active force or torque component (modelled mathematically using a Central Pattern Generator model, or CPG), a linear or nonlinear parallel elastic component, and a linear series elastic component (representing muscle tendon) (Fig. 1c). We estimated the unknown parameters of each model variation using Genetic Algorithm (GA) for predicting the wing muscle actuation torque (Fig. 1b), derived from a Computational Fluid Dynamics (CFD) simulation [19].

The derived wing-actuation model variations reveal key design traits of the hummingbird wing musculoskeletal system: 1) detailed muscle properties and functions of the two primary muscles, including active vs passive, agonist vs antagonist force generations, stress, strain, elastic energy storage, work loops, and power, and 2) spatiotemporal torque characteristics of the primary and secondary muscles in generating the 3-DoF wing motion. Together, these results show that the primary muscles do not simply flap the wing about a single stroke axis, but also actively deviate and pitch the wings with comparable torque magnitude. The capacity of pectoralis to effect these functions depends in part on a proportionally enormous bicipital crest, an evolutionarily derived trait of hummingbirds. The secondary muscles are co-activated with, and act against the primary muscles to actively constrain wing deviation and pitching. In addition, hummingbirds only use a moderate amount of elastic energy storage, as their primary muscles exhibit work loop shapes intermediate to birds and flying insects such as beetles [20], hawkmoths [21], locusts [22], and bumble bees [23]. These principles of hummingbird wing actuation may help researchers rethink and reshape the design template for agile robotic flight.

## Results

All twelve wing-actuation model variations yielded good prediction of the wing muscle actuation torque (prediction error: stroke 7.3-10.4%, deviation 7.5-14.0%, and pitching 6.6-21.0%, see Appendix), and all model variations resulted in similar contractile behaviors of PECT and SUPRA in terms of kinematics and force (Fig. 2), power and work loops (Fig. 3), and torque (Figs. 4 and 5). Model-dependent variations in the predicted muscle torque and power profiles mainly occurred in the two combined secondary muscle groups. Note that the twelve model variations were developed based on the known muscle anatomy and were used to cover the plausible musculoskeletal actuation configurations (see ‘Methods’), therefore it is expected that the regions spanned by results of the twelve model variations provide a good indication of the plausible profiles. Nonetheless, all the major results and conclusions below are independent of the model variations, while those subject to model dependency are specifically noted. Also note that every component of the defined wing-actuation model is essential since optimization of any model variation with any component removed would increase the prediction error significantly. The complete estimated model parameters are provided in Supplementary file 1; the complete results for predicted muscle functioning variables for all twelve model variations are reported in Supplementary file 2, for which median and range values are reported below.

**Fig. 2.**
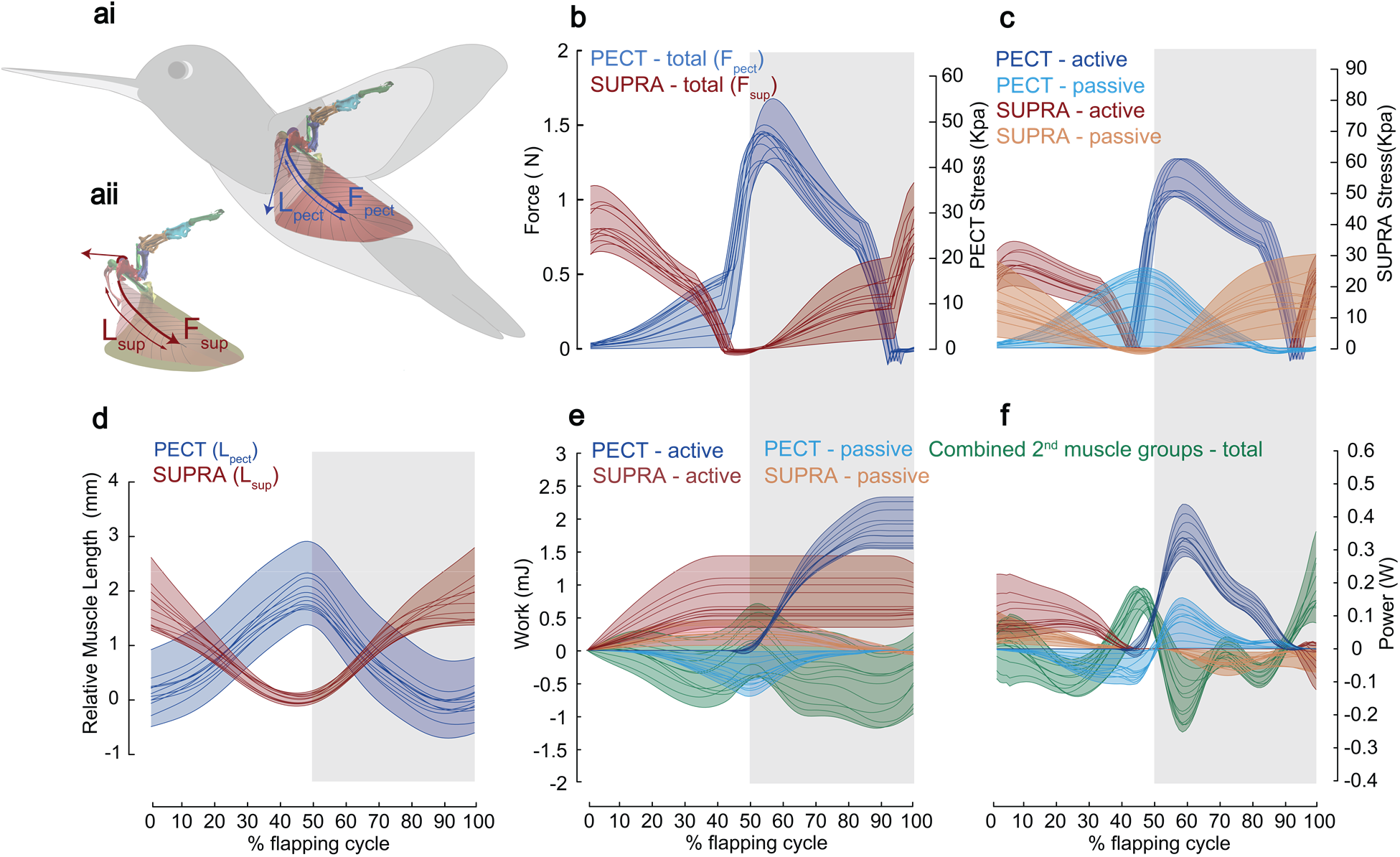
Muscle contractile behaviors, including length change, force, stress, cumulative work, and power for PECT and SUPRA, and cumulative work and power for secondary muscles. **(a)** Pictorial representation of **(i)** PECT length *L*_pect_ and force *F*_pect_ and **(ii)** SUPRA length *L*_sup_ and force *F_sup_*. **(b)** Total force and muscle stress calculated using the estimated cross-sectional area. The PECT produced cycle-averaged and peak forces of 0.54 N (median, range 0.45-0.68 N) and 1.44 N (median, range 1.25-1.68 N) (or 14-20 and 37-50 times the body weight) and the SUPRA produced cycle-averaged and peak forces of 0.34 N (median, range 0.26-0.50 N) and 0.81 N (median, range 0.65-1.11 N) (or 8-15 and 20-33 times the body weight), respectively. **(c)** Active and passive (elastic) force contributions from PECT and SUPRA. The force and stress axes are applicable to both (b) and (c). **(d)** Relative change of PECT and SUPRA length, where zero represents the mean equilibrium length. **(e)** Cumulative work and **(f)** instantaneous power by active and passive components of PECT and SUPRA, and combined secondary muscle groups over the flapping cycle, predicted by all 12 model variations. Cumulative work is set to zero at the start of the upstroke. For (b)-(f), colored shaded areas represent the regions spanned by the results for all model variations and gray shaded and unshaded areas represent downstroke and upstroke, respectively.

**Fig. 3.**
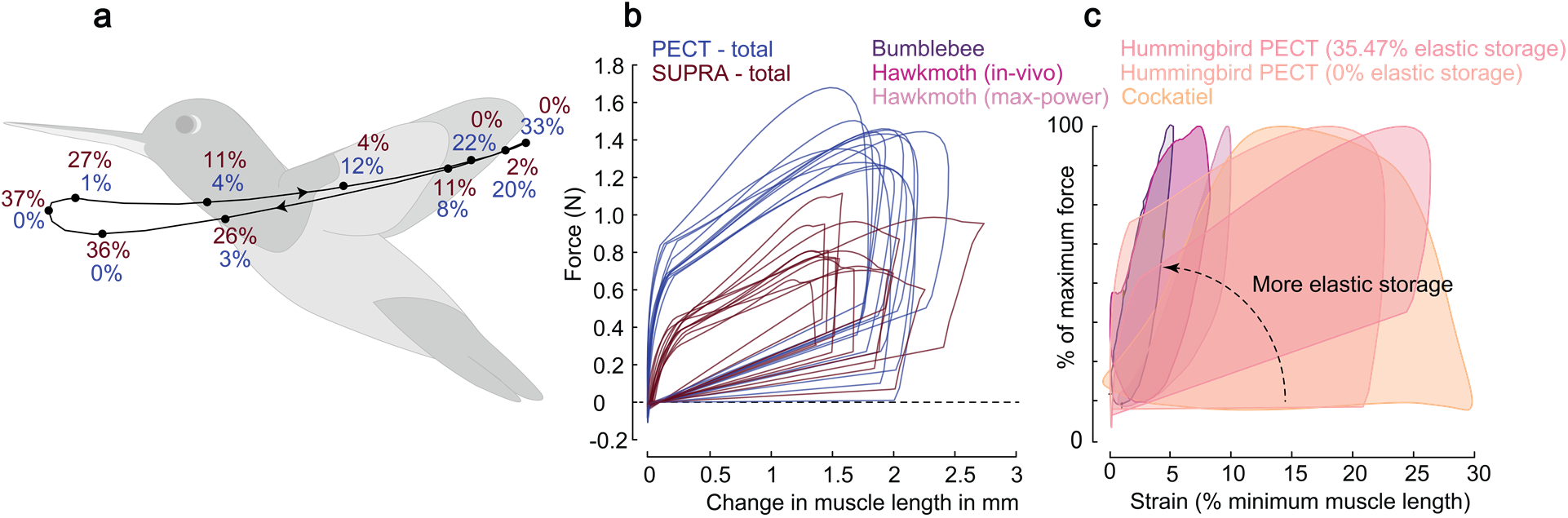
Muscle elastic energy storage and work loops for a wing flapping cycle. **(a)** Percent elastic energy storage in PECT (blue) and SUPRA (red) for wrist-centered model variations (median of the energy storage obtained for the four wrist-centered model variations, estimated at each time instant) at different instants of flapping cycle (dots on the black trace). **(b)** Work loops of PECT and SUPRA, predicted by all 12 model variations. The length shown here is relative to the minimum muscle length during the flapping cycle, which is set to zero, i.e., only change of muscle length is shown. **(c)** Work loops of PECT from two model variations with the most and the least amount of elastic storage plotted together with those of cockatiel [28], bumblebee [23], and hawkmoth (max-power and in-vivo) [21]. The work loops are scaled on the vertical axis according to the percentage of maximum force and on the horizontal axis according to the strain with respect to minimum muscle length. Note that the negative work in insects and hummingbirds during muscle lengthening indicates elastic energy storage (since active force cannot be developed at the beginning of muscle lengthening). Also note that the reproduced work loops from other species were produced using different methods. The cockatiel work loops were produced using force and strain measurements from implanted sensors while the bird was flying freely [28], the hawkmoth work loops were produced by reproducing *in-vivo* conditions on a tethered thorax and muscles, and under conditions that maximized power output [21], and the bumblebee work loop was produced for an isolated muscle fiber when it produced highest work [23].

**Fig. 4.**
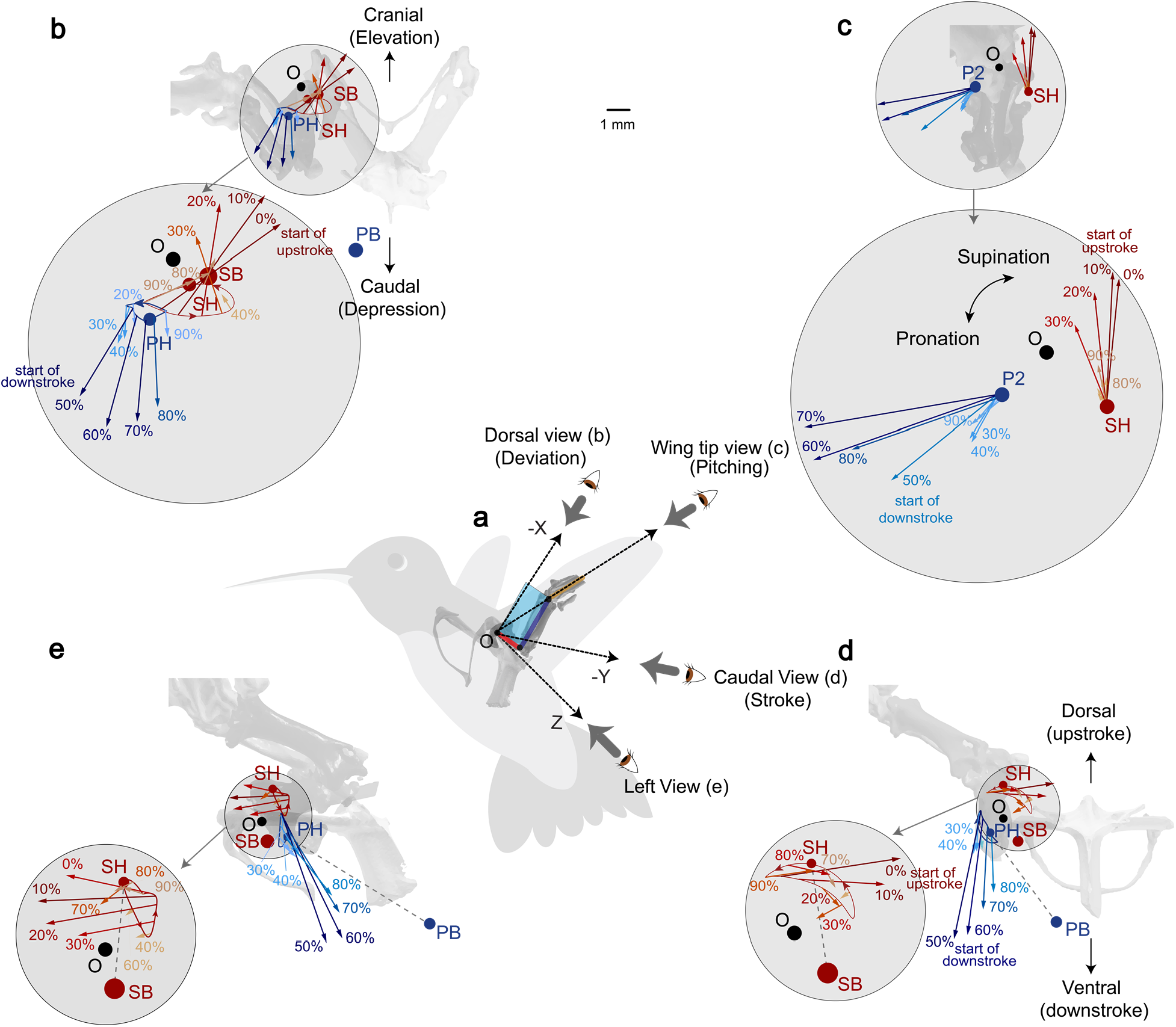
Instantaneous PECT and SUPRA force vectors over a flapping cycle. Results of wrist-centered model variations (median of the forces obtained for the four wrist-centered model variations, estimated at each time instant) are shown as an example. **(a)** Summary of four views, where PECT (blue) and SUPRA (red) force vectors are shown in **(b)** Dorsal (back) view towards the ventral body side, illustrating wing deviation torque creation (approximately); **(c)** Wing tip view in wing fixed frame from wing wrist towards the shoulder, illustrating wing pitching torque creation; **(d)** Caudal (bottom) view towards the bird’s head from tail, illustrating wing stroking torque creation; and **(e)** Left view, illustrating PECT force directions on the sagittal plane. Total ten time instants of the flapping cycle are shown (0% corresponds to the start of upstroke and 50% corresponds to the start of downstroke). Points O (black), SB (red), and PB (blue) represent the shoulder joint, insertion points of SUPRA and PECT on the body skeleton, respectively. Also shown are the trajectories of the humerus insertion points of PECT (blue) and SUPRA (red), progressing over the flapping cycle, where PH and SH represent the corresponding insertion points at the mid-downstroke instant. The dotted grey lines PH-PB and SH-SB in (d) and (e) represent the straight line connecting muscle insertion points on the wing skeleton and body skeleton during mid-downstroke. Note that the insertion point trajectories and skeletons in (b), (c), (d), and (e) (unmagnified) are scaled as per the 1 mm scale. The silhouette of the wing skeleton represents the mid-downstroke position (approximately 75% of the flapping cycle). The force vectors are scaled so that their length is proportional to their magnitude. The magnified illustrations are exactly twice the size of the unmagnified illustrations.

**Fig. 5.**
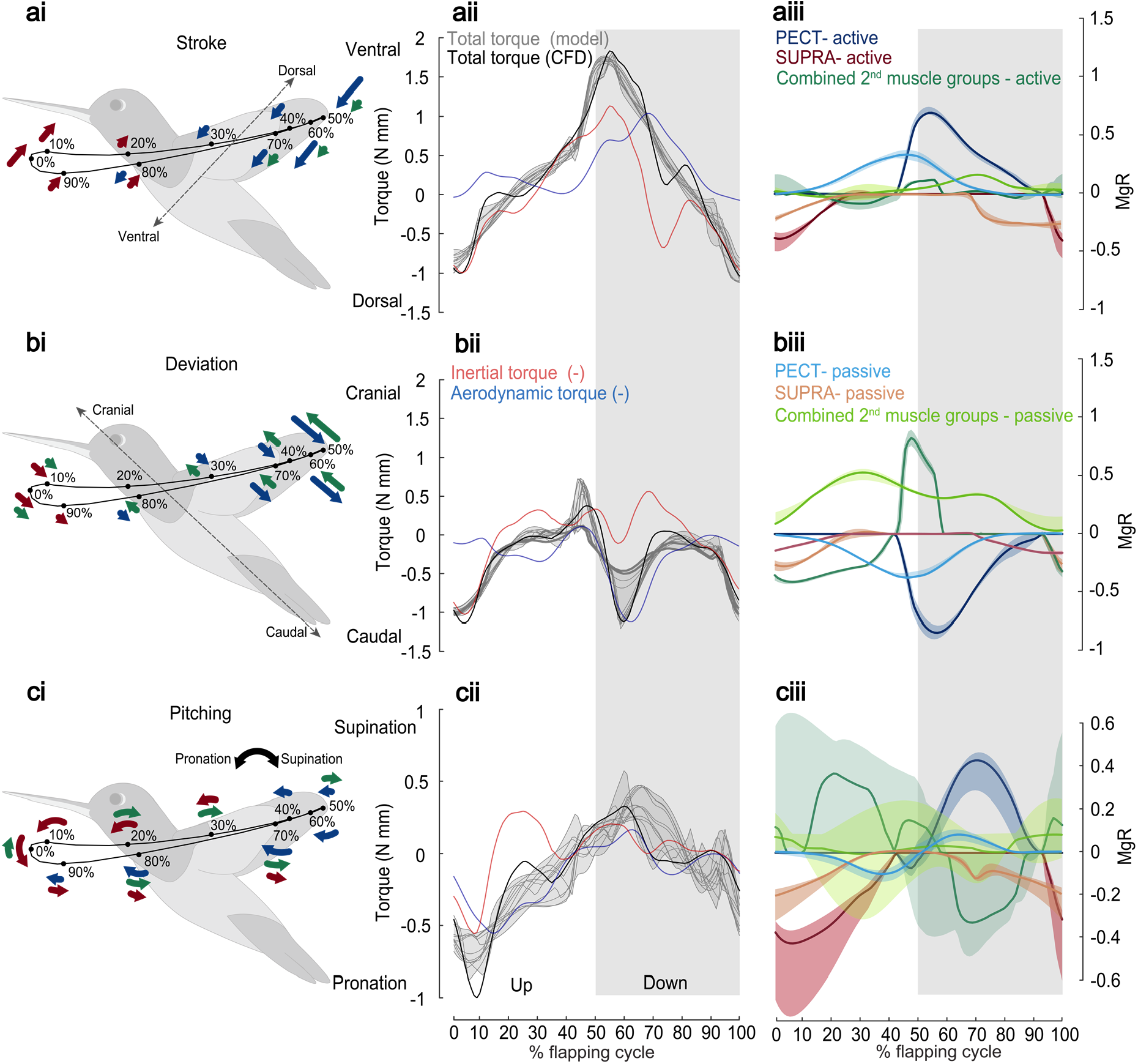
Instantaneous wing muscle actuation torque. **(i)** Pictorial representations of muscle actuation torques due to PECT (blue), SUPRA (red), and combined secondary muscles (green) for stroke **(a)**, deviation **(b)**, and pitching **(c)** axes as defined based on HUP (Fig. 1a). **(ii)** Total wing muscle actuation torque about the shoulder, predicted by our 12 model variations (solid gray) and from CFD (solid black). Also shown are the (negative) wing inertial torque (red) and (negative) aerodynamic torque (blue). **(iii)** Active and passive torque components from PECT, SUPRA, and combined secondary muscle groups, based on the wrist-centered model variations. Two torque scales are used: N-mm and the product of the body weight (Mg) and wing length (R). The solid curves in (iii) show the median torque components (median torque at each time instant estimated using the torque results for the four wrist-centered model variations) and the associated colored shaded areas represent the region spanned by torques for the four wrist-centered model variations. The gray shaded and unshaded areas represent downstroke and upstroke, respectively. The results for elbow-centered and body-centered model variations can be found in the Fig. 5 – figure supplement 1.

### Contractile behaviors of the primary muscles (PECT and SUPRA)

#### Muscle forces

The PECT of a single wing produced cycle-averaged and peak forces of 0.54 N (median, range 0.45-0.68 N) and 1.44 N (median, range 1.25-1.68 N) (or 14-20 and 37-50 times the body weight, Fig. 2b), respectively. The SUPRA produced cycle-averaged and peak forces of 0.34 N (median, range 0.26-0.50 N) and 0.81 N (median, range 0.65-1.11 N) (or 8-15 and 20-33 times the body weight, Fig. 2b), respectively. The ratio of PECT and SUPRA cycle-averaged and peak force was 1.57 (median, range 1.07-1.91) and 1.76 (median, range 1.29-2.07), respectively. These ratios are close to the ratio of their muscle cross-sectional areas [24] (0.30 cm^2^ for pectoralis and 0.21 cm^2^ for supracoracoideus, see Appendix). Both PECT and SUPRA began to produce active forces when they were still lengthening, i.e., at 8% (median, range 5.4-9.3%) of the flapping cycle period prior to the start of downstroke and upstroke (Fig. 2b), in agreement with the finding from an *in-vivo* study [25]. The contribution of elastic forces had relatively higher model dependency compared with the active forces (Fig. 2c). The ratio of the peak passive to peak active muscle force was 37% (median, range 0-51%) for PECT and 60% (median, range 12-113%) for SUPRA. This suggests that supracoracoideus likely uses higher amount of elastic energy storage than pectoralis.

#### Muscle Stress

The estimated peak muscle stress (see Appendix for calculation details) for PECT is 47.15 KPa (median, range 41.01-55.13 KPa) and for SUPRA is 38.60 KPa (median, range 31.23-53.05 KPa). The peak PECT stress obtained here is close to the pigeon peak pectoralis stress (50-58 KPa, estimated during different flight conditions) and the peak SUPRA stress is lower than the pigeon peak supracoracoideus stress (85-125 KPa, estimated during different flight conditions) [26].

#### Change of muscle effective length and strain

The effective muscle lengths of PECT and SUPRA were calculated as the lengths of the curved arcs (Fig. 2a) along which the muscles lie (see ‘Methods’). We assumed that the muscles were relaxed at the end of their respective half-strokes and were maximally stretched at the start of their respective half-strokes. The estimated PECT length change (Fig. 2d) resulted in muscle strain ranging from 18% to 24% with respect to the estimated mean PECT length. These strain values are higher than that measured for pectoralis in rufous hummingbird (11%) [25], comparable to those obtained *in vivo* in zebra finch (17-21%) [27], but significantly lower than those reported in other birds (29-44% during take-off, ascending, descending, and level flight for cockatiel [28,29], mallard [30], and pigeons [31,32]). The estimated SUPRA length changes appear comparable or slightly lower than those of PECT (Fig. 2d), however, we could not estimate the SUPRA strain as its effective origin on the sternum could not be estimated.

#### Mechanical work

The cumulative work done over a complete flapping cycle was 1.78 mJ (median, range 1.55-2.33 mJ) by PECT and 0.58 mJ (median, range 0.35-1.27 mJ) by SUPRA (Fig. 2e). Active, non-elastic force components produced positive power consistently over the flapping cycle (with only minor instantaneous power absorption) (Fig. 2f), resulting in positive net work output. On the other hand, passive, elastic forces stored and released energy, and produced zero net work. Specifically, the PECT elastic component stored energy (negative power) at the later part of upstroke and released it at the beginning of downstroke (Fig. 2f and 3a), as speculated previously for ruby-throated [33,34], rufous [35], and broad-tailed [35] hummingbirds. Similarly, the SUPRA elastic component stored energy in the later part of downstroke and released it at the beginning of upstroke. Notably, elastic energy storage plays a crucial role in improving muscle efficiency, as previously reported in hummingbirds [33–35] and insects [36,37]. The maximal muscle elastic energy stored, as percentage of total cumulative work, was 26.43% (median, range 0%-35.47%) for PECT and 43.66% (median, range 7.46%-75.96%) for SUPRA. These median values are considerably higher than those in pigeons (pectoralis: 18%, supracoracoideus: 14%) [26] but comparable or lower than many insects (e.g., locusts [38]: 50%, bumblebees [39]: 30%). Moreover, our results underscore the non-negligible work contribution from supracoracoideus in contrast to other birds [40], due to the more active upstroke in hummingbirds. Together, these results further support the argument that hummingbirds have been evolutionarily converging towards insects, as their primary muscles use proportionally more elastic energy storage than those of other birds [41,42].

The work done by combined secondary muscle groups showed more significant model-dependent variations, from absorbing large negative work (−0.97 mJ) to producing small positive work (0.27 mJ), although all model variations agreed that the secondary muscles together produce substantially less net positive work than each of PECT and SUPRA (Fig. 2e). This is consistent with the small size of these muscles, rendering them incapable of providing large positive work, despite producing large torque (see below). Interestingly, the peak instantaneous power output by combined secondary muscle groups could be comparable to that of PECT (Fig. 2f).

#### Mass-specific power

The estimated muscle mass-specific power (see Appendix for calculation details) is 377 W kg^−1^ for PECT (median, range 342-491 W kg^−1^) and 408 W kg^−1^ for SUPRA (median, range 260-837 W kg^−1^). Note that these estimates include wing inertial power with imperfect muscle elastic energy storage, and are therefore higher than those reported for other species of hovering hummingbirds (185-228 W kg^−1^) [43] that assumed perfect muscle elastic energy storage.

#### Work loops

The work loops [24,44] of PECT and SUPRA (Fig. 3b) can be obtained based on their respective force and length profiles. The area underneath a work loop curve represents the muscle work, which is positive during shortening (muscle active and elastic components output energy) and negative during muscle lengthening (muscle elastic component absorbs energy). Accordingly, the area enclosed by a work loop corresponds to cumulative work over a complete cycle. The PECT and SUPRA work loops obtained here are intermediate among those of other birds (Fig. 3c) (i.e., pigeon [26], budgerigar [45], cockatiel [28,29], and zebra finches [46]), and insects (i.e., locust [22], bumblebee [23], hawkmoth [21], and beetle [20]). The peak force is produced near the middle of the muscle shortening phase in other birds [26,28,29,45,46] and near the end of the muscle lengthening phase in insects [20–23]. Hummingbird pectoralis, instead, starts to produce force towards the end of the muscle lengthening phase with a short active lenthening, followed rapidly by the force peak in the beginning of muscle shortening phase.

### PECT and SUPRA instantaneous force vectoring

In all hypothesized wing-actuation model variations, both PECT and SUPRA force directions were allowed to rotate linearly (with respect to the instantaneous straight-line PH-PB connecting muscle insertion points on the wing skeleton and body skeleton, Fig. 4d-e) with the progression in the flapping cycle, in accordance with the anatomical features. Results show that at the beginning of downstroke, PECT force direction was oriented ventrally by 46° (median, range 38° to 48°) from the straight-line PH-PB (Fig. 4d-e, see Appendix and Movies S1, S2, and S3). During downstroke, PECT force rotated dorsally and became almost aligned with PH-PB towards the end. Similarly, at the beginning of upstroke, SUPRA force was oriented dorsally by 57° (median, range 39°-88°) from SH-SB (Fig. 4d-e) and became more aligned with SH-SB (median 9°, range 0°-44°) at the end of upstroke. Overall, the vectoring of PECT and SUPRA forces show highly elaborated spatiotemporal patterns (Fig. 4), which are more prominent than those observed in other birds such as pigeons [31]. As a result, they lead to highly three-dimensional wing actuation torque described below.

### Wing muscle actuation torque characteristics: primary vs secondary muscle groups and active vs passive muscle torques

The wing muscle actuation torque is characterized here about stroke, deviation, and pitching axes defined on HUP (which can be considered as the rigid, proximal part of the wing, Fig.1a), since all the muscles that transfer force from a hummingbird’s body to its wing insert only on the bones that lie entirely on HUP. For clarity, only results from wrist-centered model variations are shown in Fig. 5, while those from elbow-centered and body-centered model variations are shown in Fig. 5 – figure supplement 1.

#### Stroke

The wing stroking torque was mainly generated by PECT and SUPRA (Fig. 5a), with negligible contribution from combined secondary muscle groups. Both PECT and SUPRA were activated prior to the start of their respective half strokes and subsequently provided peak stroking torque near the start of half strokes (Fig. 5ai). The peak stroking torque of PECT was estimated 1.51 (median, range 1.03 to 1.90) times higher than those of SUPRA (Fig. 5aiii), while this range includes the ratio of 1.5 reported for pigeons [32]. The peak passive stroking torque produced by these two muscles were comparable (Fig. 5aiii).

#### Deviation

The wing deviation torque mainly counterbalanced the elevation torque due to aerodynamic lift at the beginning of downstroke (blue, Fig. 5bii), and overcame wing inertia near the beginning of the upstroke (red, Fig. 5bii). The wing deviation torque was primarily generated by PECT at the beginning of downstroke and by SUPRA at the beginning of upstroke, both in the caudal (depression) direction (Fig. 5b). The secondary muscle groups, however, produced elastic torque that acted against the depression torque from PECT and SUPRA over the entire flapping cycle (Fig. 5biii). In addition, they also produced a large active elevation torque during the downstroke against the PECT depression torque. The active elevation torque from the secondary muscle groups showed some variation among body-centered, elbow-centered, and wrist-centered model variations; however, in general, they tended to counteract against the two primary muscles, especially during downstroke, to create a controlled-tightening effect.

#### Pitching

The wing muscles functioned as “active stoppers” to resist the passive wing pitching (Fig. 5c), generated by both aerodynamic and inertial effects in the first half of each half stroke (Fig. 5cii). Specifically, SUPRA acted to pronate the wing in ventral stroke reversal when the wing was being supinated passively, and PECT and secondary muscles acted primarily to supinate the wing in the beginning of downstroke when the wing was being pronated (Fig. 5cii). Similar to deviation, there was consistent counteraction between torques due to primary and combined secondary muscle groups (Fig. 5ci and ciii) near the middle of each half-stroke, despite more significant model-dependent variations (Fig. 5 – figure supplement 1c). Notably, for elbow-centered model variations, the active pitching torque due to secondary muscles was nearly zero throughout the flapping cycle while the active pitching torque due to primary muscles was lower (Fig. 5 – figure supplement 1c).

Note that although the deviation and pitching torques generated by secondary muscles were comparable to the stroking torque in magnitude, their forces are expected to be significantly lower than those of primary muscles, because they use substantially larger moment arms (Fig. A3c, Appendix). Secondly, for both PECT and SUPRA, we found that if the force vector was directed linearly from its insertion point on the humerus to that on the body skeleton (i.e., without redirection offered by the curved bicipital crest), the relative contribution towards stroking would decrease, while those towards deviation and pitching would increase significantly.

### Near co-activation of primary and secondary muscle groups

The counteracting action described above suggests that the primary and secondary flight muscles are nearly co-activated. This is further validated by examining the onset timings optimized for model variations that assumed independent activation of different muscle groups (Fig. 1d). Among these model variations, we found relatively small differences in the onset timings of active force or torque between the corresponding muscles from the three muscle groups. The maximum phase difference was approximately 5% (of flapping cycle period) between the activation of PECT and one of the DMG muscles, and approximately 8% between the activation of PECT and one of the PiMG muscles (Supplementary file 1i).

## Discussion

### Primary flight muscles function as diverse effectors that rotate the 3DoF shoulder, which is actively tightened by secondary muscles in deviation and pitching

The wing muscle actuation torque characteristics (Fig. 5) reveal that the primary muscles, i.e., hummingbirds’ “flight engine”, do not simply ‘flap’ the wing along a single DoF, as the wing motion *per se* might appear to be; instead, they actuate the wing via an approximate ‘ball-and-socket’ joint, generating torque of comparable magnitude in all three wing axes of stroke, deviation, and pitching. The deviation and pitching are actively constrained by sizable counteracting torque from secondary muscles, resulting in a controlled tightening effect and relatively small wing excursion (Fig. 6). Only in the stroke direction, however, the wing is allowed to rotate largely unconstrained so that there is a large excursion and muscle power output, giving an impression of approximately 1-DoF wing motion to an observer (Fig. 6).

**Fig. 6.**
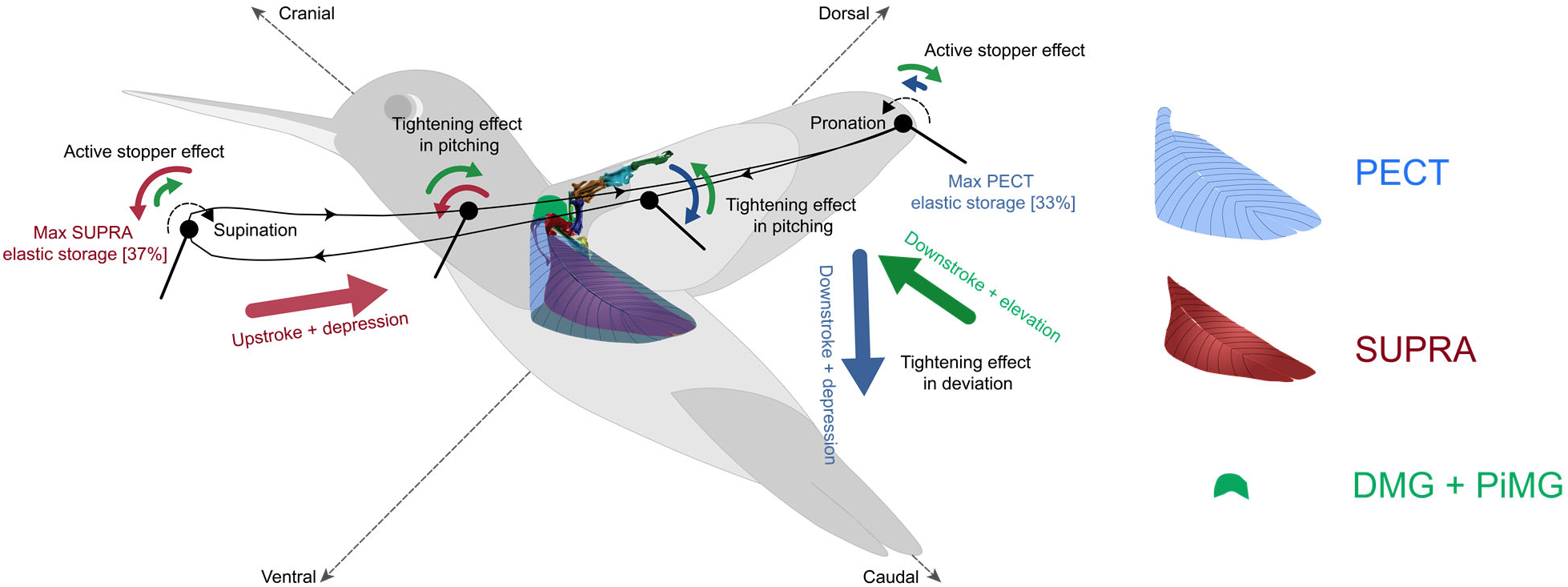
Summary of hummingbird wing musculoskeletal actuation principles. The pectoralis (blue) and supracoracoideus (red) (referred to as PECT and SUPRA in the wing-actuation model) function as diverse effectors, generating actuation torque in all stroke, deviation, and pitching directions with comparable magnitude. Pectoralis generates wing downstroke and depression (i.e., pull the wing ventrally and caudally) and supracoracoideus generates wing upstroke and depression (i.e., pull the wing dorsally and caudally). Pectoralis, supracoracoideus, and the combined secondary muscle group also function as ‘active stoppers’ for pitching, acting against the pronation or supination due to the aerodynamic and inertial effects. The combined secondary muscles (green) create controlled tightening effects in deviation and pitching by counteracting the primary muscles. The pectoralis and supracoracoideus muscles can also store maximal elastic energy of 37% and 33% (of the total energy output, these percentage values are the median of the maximum energy storage obtained for the four wrist-centered model variations), respectively. See Movie S1 for illustration of these principles alongside wing skeleton movement.

One potential benefit of such tightening, in the axes approximately orthogonal to the primary stroke axis, could be acquiring high wing agility and stability along deviation and pitching, which translate to higher control authority of flight dynamics. This is similar to the effects of the non-propulsive, counteracting lateral forces (orthogonal to the direction of locomotion) in legged locomotion [47]. Comparing wing deviation and pitching, deviation is likely a more agile axis for wing motion modulation and therefore is likely recruited more extensively in flight control. This is because wing pitching dynamics often involve more complex, highly nonlinear wing camber and twisting (in both feathers and wrist joints), both of which could render a more complex “plant” [48] for control. Changes in wing deviation, however, are less dependent on wing structural dynamics; they create not only fore/aft and lateral tilting of the wing stroke plane for force vectoring, but also modulate the wing angle of attack. Observations of hummingbird escape maneuvers [1,43] seem to confirm these speculations, in that wing deviation exhibited the largest changes, and is indeed the strongest contributor for generating body pitch (via fore/aft tilting of the stroke plane) and roll torque (via lateral tilting of the stroke plane), compared with changes in stroking and pitching. We predict that by changing the activation of secondary muscles, and with a relatively small force and power output, hummingbirds can modulate wing deviation to produce subtle, less intensive maneuvers [49]. On the other hand, hummingbirds can also produce large-magnitude, more intensive maneuvers, by changing the activation of primary muscles that lead to concurrent increase in stroke velocity and spatiotemporal modulation in deviation motion [1,43] (see next section). It is also worth noting that insects with advanced flight capacity and smaller body size than hummingbirds, such as flies and bees, do not seem to have the capability of modulating wing deviation and pitching directly using their power muscles [50]. Instead they rely mainly on smaller steering muscles for generating higher-order perturbations to the primary wing motion created by thoracic oscillation, which is driven by asynchronous power muscles [51,52]. The elevated capacity in wing deviation and pitching modulation might explain why hummingbirds can vector their aerodynamic forces to a much greater extent and do not conform to the helicopter model as much as do insects [1,53].

### The adaptation in humerus enables pectoralis force vectoring that promotes both hovering and flight maneuverability

The humerus of hummingbirds has a morphology and function that is substantially different from those of other birds [54,55]. It was previously found that a high degree of long-axis rotation enables primary muscles with low-strain to actuate large-amplitude wing motion [8]. Our results suggest that a significantly larger protrusion in the humerus (i.e., bicipital crest, Fig. A2) magnifies this effect while also facilitating optimal orientation of pectoralis force. First, the pectoralis force application point on the humerus is shifted away from the long-axis due to its insertion on the proportionally enormous bicipital crest. This increases the moment arm for both wing stroking and pitching torque. Second, the shape of bicipital crest appears to have evolved for properly orienting the pectoralis force direction. The hummingbird pectoralis can pull the humerus more caudally (tailward) and dorsally compared with, for example, pigeons [26,56], thereby directing the pectoralis force more tailward for larger wing depression (deviation) torque (Movie S3). This appears to be enabled by a more caudal orientation of the pectoralis muscle fibers or a more caudal insertion of the pectoralis on the sternum. Such anatomy is necessary to balance the larger lift-induced wing elevation torque, as hummingbirds hover with an upright body posture. On the other hand, the bicipital crest on humerus, where the pectoralis aponeurosis wraps around, prevents an excessive caudal redirection of the force (Fig. 4e), ensuring proper distribution of the force into creating both stroking and deviation torque (a direct connection between the pectoralis insertion points on the humerus and sternum would make the force directed mostly caudally). As a result, the pectoralis force generates concurrent downstroke, depression, and supination torque in stroke, deviation, and pitching directions, respectively. Such coupled torque generation is remarkably consistent with the wing kinematics modulation observed in the initiation phase of the hummingbird escape maneuver [1]; the bird simultaneously increases its stroke amplitude and frequency (increases total aerodynamic force), tilts its stroke plane backwards by depressing its wing dorsally (directs the aerodynamic force backward), and uses larger supination during downstroke (increases the angle of attack and therefore backward force or maintains a proper angle of attack when wing loading increases due to higher stroke velocity). This suggests that via a single downstroke, the hummingbird wing musculoskeletal system can use the elevated pectoralis force to generate changes in wing stroke, deviation, and pitching that are coordinated to generate large body backward force and pitching torque, likely promoting maneuverability. Therefore, while using kinematically “rigid” wings without the flexion exhibited during upstroke in other species [57], hummingbirds seem to have more sophisticated torque control around the shoulder than other birds, thanks to the adaptations in their humerus. We hypothesize that the evolutionary enlargement of the bicipital crest was a key process leading to the unique capacity of pectoralis in hummingbirds.

### Hummingbirds use moderate elastic energy storage to save wing inertial power at high wingbeat frequency, while possibly avoiding full mechanical resonance to retain direct motor control for high wing agility

The cumulative work profiles (Fig. 2e) and the work loop shapes (Fig. 3b-c) suggest that pectoralis and supracoracoideus of hummingbirds use higher amount of elastic energy storage than other birds (such as pigeons [26], budgerigar [45], and cockatiels [28,29]), but less than many insects with comparable or smaller body size (such as locusts [38] and bumblebees [39]). In those insects, muscular elastic energy storage (area beneath the workloop, Fig. 3c) is comparable or higher than the energy dissipated to overcome aerodynamic drag (area enclosed by the workloop); while for larger birds, majority of the muscular energy is spent to overcome aerodynamic drag with negligible elastic energy storage. The predicted work loops of PECT resemble more to those of larger birds but are shifted towards those of insects since hummingbirds use moderate elastic energy storage, the amount of which remains considerably lower than the energy dissipated. This trend is also evident when comparing the timings of peak PECT (or power muscle) force (Fig. 3c), which are attained near mid-downstroke in other birds (such as pigeon and cockatiel [26,28]), shortly after the beginning of downstroke in hummingbirds, and at the maximum muscle length (e.g., at the beginning of the downstroke) in insects, especially those who use asynchronous flight muscles, who likely use the greatest amount of elastic energy storage and mechanical resonance [58].

This result not only further supports that hummingbirds have evolved towards insect flight, but may also reflect a tradeoff between muscle efficiency and direct motor control capacity of primary muscles. High elastic energy storage improves muscle efficiency for hovering at high wingbeat frequency and also alleviates the demand for fast motor control since the flapping frequency is mainly determined by that of mechanical resonance [36]. However, these benefits are at the cost of lower capacity for active, direct muscular control of wing motion, due to strong mechanical resonance, which can only be modulated indirectly, e.g., by variable stiffness [59,60], or to a limited amount by steering muscles [61]. Therefore, this tradeoff could possibly influence the convergent evolution of hummingbirds towards insects. While hummingbirds have evolved towards hovering capacity with high wingbeat frequency and more symmetric flapping, they need to use *more* muscle elastic storage to overcome larger amount of wing inertial power, but *less* muscle elastic storage to retain direct muscular control of wing motion, and the solution seems to be in the middle ground, i.e., having a moderate amount of elastic storage. Together, the unique actuation capacity and the work loops of hummingbird primary muscles might also explain the distinct scaling rules in their wing morphology and frequency, when compared with other birds and flying insects, as previously observed by C. Greenwalt [62].

### Principles of hummingbird wing actuation do not seem followed by the existing designs of hovering-capable, hummingbird-inspired robots

In the past two decades, flapping flight in hovering-capable animals, such as hummingbirds and insects, has been perceived as a novel form of flight that can be emulated in micro aerial vehicles for improved flight stability and maneuverability. However, despite the success in developing flapping-wing robots with sustained hovering and basic maneuvering capacity [5,7,12–18], these robots do not outperform rotary-wing micro aerial vehicles in terms of maneuverability and hovering efficiency.

So what is missing? If we examine the existing designs of hovering-capable, flapping-wing robots with wingspan similar to that of hummingbirds, and try to summarize the principles they follow, we can find the following – first, the power actuators (e.g., electric motors [7,12–14,16,17] or electromagnetic actuators [18]) only drive a single axis wing motion [7,12–18], i.e., flapping or stroking, while wing pitching is either passive [7,16–18] or independently modulated via twisting or slacking the wing surface [12–15]; second, wing deviation is not used in the flight control and it is entirely hard constrained; and third, half of the designs tend to maximize elastic energy storage [17] or promote a resonant actuation system [16–18] to improve efficiency. These principles do not follow those of the hummingbird wing actuation – first, hummingbird power muscles actuate all 3DoF wing motion in a coordinated fashion to promote maneuverability, while actively constraining wing pitching and deviation; second, wing deviation is likely the most agile axis of wing motion control and used most extensively in escape maneuvers [1,43]; and third, hummingbirds only use a moderate amount of elastic energy storage, likely trading off muscle efficiency and flight control authority. In short, hummingbirds do not simply flap their wings as most of their robot mimics do, as the relatively simplistic designs adopted by the latter do not capture the essential design traits of hummingbird flight.

These principles and their implications on robotics also need to be interpreted with the caution of size-dependency, as they may not directly apply to smaller fliers, such as flies, bees, or robots of similar size. It is known that smaller body size poses more stringent constraints on wing actuation designs [63], however, offers inherently higher rotational maneuverability [43,64], which for smaller fliers, is mainly limited by sensorimotor control, rather than wing actuation [2]. Therefore, for smaller flying insects, it is plausible that a high-frequency, primarily single-axis wing flapping generated by a strongly coupled thorax-wing oscillation, is size-appropriate. While such thorax-wing oscillation is driven by asynchronous flight muscles, higher-order and small wing kinematic changes are effected and controlled by steering muscles [52,63]. Hummingbirds, on the other hand, use their musculoskeletal wing actuation to generate larger amounts of wing kinematic changes and force vectoring, thereby attaining comparable rotational maneuverability to their smaller insect counterparts [2], and also higher linear maneuverability due to their larger body size [43]. In sum, body size could be a critical factor when developing templates for flapping-wing robots inspired by hummingbirds or flying insects.

So how could these understandings help advance agile robotic flight? These principles do not directly translate into specific design guidelines or templates, and robot design and fabrication are subject to a myriad of technical challenges in actuators, materials, onboard electronics, power, sensors, and constraints in size and weight. Nonetheless, it is conceivable that these principles could help researchers, at least start to rethink and reshape the current design templates, hopefully leading to renewed attempts for designing flapping-wing robots that truly leverage the high control authority provided by flapping flight [65] with large-amplitude, high-frequency, and controlled motion of their aerodynamic surfaces.

## Methods

The hummingbird wing musculoskeletal system can be described as a 3-link, 7-DoF system (Fig. 1a) actuated by at least 40 muscles [9]. The torque transfer from the bird’s body to the wing musculoskeletal system however is enabled by only roughly 14 muscles that connect the body skeleton and the first two links of the system, i.e., humerus and radius/ulna (which define the HUP, Fig. 1a). Our musculoskeletal wing-actuation model considers the actuation of the wing skeleton, by a Primary Muscle Group (PMG), including pectoralis and supracoracoideus (referred to as PECT and SUPRA in the wing-actuation model) that inserts on the humerus, and two combined secondary muscle groups, i.e., Pitching Muscle Group (PiMG) and Deviation Muscle Group (DMG) that insert on both the humerus and radius/ulna.

The process of developing the wing-actuation model and its variations involved three steps. First, we developed a hypothetical model and its variations with unknown parameters, based on the hummingbird skeletal and muscular anatomy and kinematics. We synthesized data from multiple sources and also conducted additional dissections, for finding the dimensions, orientation, function, and insertion points or regions of different muscles throughout a flapping cycle. We hypothesized twelve model variations to cover all conceivable possibilities of the wing musculoskeletal system functioning. Second, we estimated the wing muscle actuation torque using the aerodynamic and inertial force data from the CFD simulations conducted by Song et. al. [19]. Finally, we optimized the unknown parameters of each model variation to minimize the difference between model-predicted and CFD-estimated wing actuation torque. The following sections describe the details of these steps.

### Hypothetical musculoskeletal wing-actuation model and its variations

To develop the model and its variations, we integrated the wing skeletal kinematic data [8] with the hummingbird anatomical data [9,10]. First, we used the X-ray marker data and the μCT scan of a hovering ruby-throated hummingbird [8], for dimensions and orientation of the wing skeleton. The hummingbird wing skeleton is approximately a 3-link, 7-DoF system, where the shoulder and wrist joints can be modelled as 3-DoF ball-and-socket joints and the elbow joint can be modelled as a 1-DoF hinge joint (Fig. 1a). We calculated the rotation angles for all the joints at each instant of the flapping cycle based on the skeletal kinematics [8] (Fig. 1a), referenced with respect to the mid-downstroke position (see Appendix for calculation details). Second, we examined the previously reported attachment points or regions, and functions of the known muscles connecting the bird’s body skeleton to the wing skeleton (Fig. A1, Appendix and Supplementary file 3). In this effort, we synthesized anatomical data from two previous studies [9,10], which include a detailed and comprehensive account of the muscle anatomy of purple-throated Carib and Anna’s hummingbirds, as well as the data from the other birds (see Appendix) where the data for hummingbirds was not available. Third, we also conducted dissection on a calliope hummingbird to find the exact attachment region for pectoralis, supracoracoideus, and M. scapulohumeralis caudalis (referred to as SHCA in the wing-actuation model), and their possible orientation (Fig. A2 and A3, Appendix).

The developed wing-actuation model comprises of a primary muscle group, which includes the antagonistic PECT and SUPRA, and two secondary muscle groups, which are used to model the combined torque-generation function of all other muscles. Note that each muscle group consisted of two muscles (for PMG, they are PECT and SUPRA), which provide torque in same or opposing directions. The followings describe the mathematical formulation of an individual muscle model, the details behind each muscle or muscle group model, and other relevant modeling details.

#### Mathematical model of an individual muscle

The generation of muscle force or torque of an individual muscle was modelled using a generic three-component model, which has been widely adapted [66]. The muscle model is comprised of an active forcing component, a series spring, and a parallel spring (Fig. 1c). The model can be described mathematically by Eq. 1, 2, or 3 for linear, cubic, or combined linear and cubic springs, respectively,

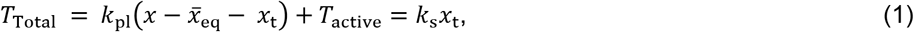

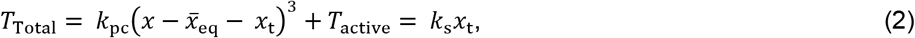

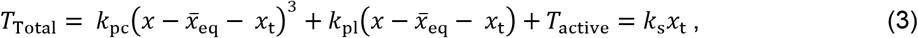

where *x_t_* is the change in tendon length, *x* is the total muscle length, 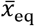 is the muscle equilibrium position, *T*_active_ represents the active force component, and *k*_pl_, *k*_pc_, and *k*_s_ are the spring constants for linear parallel spring, cubic parallel spring, and series spring, respectively. The series spring was assumed to be linear since it was previously found that the tendon elastic characteristics are linear [67]. The active force of the muscle (*T*_active_) was modelled using a central pattern generator (CPG) model, since its output closely mimics the actual muscle active force profile (see Appendix).

In addition, we provided the provision for the active force to be initiated at any instant of the flapping cycle by including additional parameters *ϕ, ϕ*_DMG_ and *ϕ*_PiMG_, known as the PMG, DMG and PiMG active force onset phases (see Appendix).

##### Details of individual muscle or muscle group model

###### Primary Muscle Group (PMG); PECT (Pectoralis) and SUPRA (Supracoracoideus)

We first estimated the effective pectoralis insertion (or force application) point on the humerus. Through dissection, we found that pectoralis muscle fibers and aponeurosis are connected to the humerus via a narrow, long region (yellow region in Fig. A2), instead of a single point. Therefore, this gave the possibility that the effective point of muscle force application may not be fixed and can be time-varying, i.e., changes throughout a flapping cycle. However, the results of model optimization permitting this change resulted in an insertion point at the bottommost connection point (PH_2_, Fig. A2 and A3) or did not improve costs significantly, therefore, we assumed that PECT attaches at point PH_2_ for the final optimization (see Appendix). The supracoracoideus effectively attaches at a single attachment point as found earlier [9] (Fig. A1a) and through dissections.

We then considered the possibility of independent activation of different muscle fibers. It was previously found that for pigeons, different regions of pectoralis muscle are activated almost concurrently [31,56]. Since pectoralis and supracoracoideus of hummingbirds are activated by a smaller number of spikes, with significantly higher wingbeat frequency [68], we assumed that different muscle fibers are activated concurrently via a single spike. We also assumed that the force due to each muscle fiber spans an entire half stroke (see Appendix).

We then estimated the direction of the total pectoralis force. Since muscle fibers in pectoralis are evenly distributed on both sides of the aponeurosis (Fig. 1ci), it was assumed that their combined force effectively acts tangent to the aponeurosis at the effective humerus insertion point (Fig. A2, Appendix). In addition, the aponeurosis (and the overall muscle) wraps around the bicipital crest (elongated protrusion) on the humerus (See Fig. 1ci, A2, and A3) and the overall tendon and muscle were assumed to have non-zero bending modulus. Therefore, a non-zero angle was included for orienting the force vector with respect to the vector PH-PB, i.e., the nominal direction connecting PECT insertion points on the body and humerus (see Fig. 4d). We used two variables (*θ_pz_* and *θ_py_*, see Appendix) to represent the overall 3D rotation of PECT force vector from its nominal direction (PH-PB, Fig. 4d). Similar to PECT, SUPRA force vector was assumed to lie tangent to the muscle tendon at the humerus insertion point (red vector in Fig. 1ci) and that the tendon has non-zero bending. The final force vector was found by rotating the nominal vector SH-SB (Fig. 4d) in the tendon plane by *θ_s_* (see Fig. A3 and Appendix). Both the muscles were modeled using arcs along their aponeurosis connecting their two insertion points on the body and humerus, being tangent to their muscle force direction at the humerus. Accordingly, the muscle length was then calculated as the arc length (see Appendix), which was also used as a length measure for calculating the passive muscle force due to the parallel spring. Note that the length of muscle fascicles is proportional to this estimated muscle length, assuming that the pennation angle of the fascicles with respect to aponeurosis remains constant throughout the flapping cycle.

We assumed that both PECT and SUPRA are in relaxed state near their minimum lengths, i.e., near the end of downstroke and the end of upstroke. We therefore allowed the equilibrium position of PECT and SUPRA to be a maximum of 0.1 mm higher than their minimum muscle lengths.

We also included SHCA separately in our wing-actuation model, however, it did not improve our model prediction. Therefore, SHCA was not included explicitly in the final model variations (see Appendix).

###### Deviation Muscle group (DMG)

The DMG modeled the torque generation from combined secondary muscles (other than PMG muscles) that was hypothesized to be acting along either the HUP perpendicular axis (DMG_h_ or DMG_w_, Fig. 1cii) or the bird’s ventral axis (DMG_b_, Fig. 1ciii) for different model variations (Fig. 1d). The muscle length was approximated either by the instantaneous length between points SeB and SeR (L_WL_, Fig. 1d and Fig. A3c) or by the Euler deviation angle for wrist or elbow centered HUP (θ_w_ or θ_e_, Fig. 1d). These length measures were hypothesized to proportionally approximate effective muscle length of combined secondary muscles.

The torque magnitude due to DMG was modelled similarly to the force magnitude due to force-generating primary muscle group; however, unlike PECT and SUPRA, the active torque due to each muscle could span any percentage of the half stroke, which is referred to as muscle active period (see Appendix), and was set as a model parameter for optimization. We also provided the provision for the active torques due to the two muscles to act along same or antagonistic directions by including binary parameters for optimization (see Appendix).

###### Pitching Muscle group (PiMG)

The PiMG modeled the torque generation from secondary muscles (other than PMG muscles) that was hypothesized to be acting along either of the HUP long-axis (PiMG_w_, Fig. 1cii), humerus long-axis (PiMG_h_, Fig. 1cii), or the perpendicular to the body skeleton (PiMG_b_, Fig. 1ciii). The muscle length was approximated by either of L_WL_, the long-axis rotation angle of the humerus (L_HR_), or the Euler pitching angles for wrist or elbow centered HUP (Ψ_W_ or Ψ_E_, Fig. 1d). The torque magnitude due to PiMG was modelled similar to DMG, and the direction of the two active torques were also optimized using binary parameters.

###### Phase relationship of active force or torque onset times among different muscle groups

The phase difference between active force or torque onset times of the two muscles within each muscle group was fixed at 50% of the flapping cycle period (see Appendix for details). However, inter muscle group onset times were allowed to be identical or independent. Specifically, the active torque onset of one muscle from DMG or PiMG can be identical or independent to that of PECT and the active torque onset of the other muscle can be identical or independent to that of SUPRA (Fig. 1d).

###### Twelve model variations

Collectively, there were two model variations depending on whether the active force or torque onset times are independent or identical between primary and secondary muscle groups, three model variations depending on the actuation axes of DMG and PiMG, and another two model variations depending on the effective muscle length estimates. In total, this yielded a total of 12 model variations (Fig. 1d).

###### Elastic characteristics of shoulder joint for model variations 5, 6, 10, and 12

For these model variations where DMG and PiMG act along X and Y axes (Fig. A3a), a separate elastic component representing the joint elasticity along the stroke direction (SSPR_b_, Fig. 1ciii) was included since DMG and PiMG in these particular models do not provide elasticity along stroke direction while the shoulder joint is likely to possess elasticity along the stroke direction due to the elastic characteristics of muscles and tendons other than PECT and SUPRA.

### Estimation of wing muscle actuation torque based on CFD data

The total muscle actuation torque transferred from the bird’s body to its wing was estimated based on sum of the wing aerodynamic forces and inertial forces, following an inverse dynamics approach. A brief overview is provided below and the full details of this estimation can be found in the Appendix.

We first calculated the wing inertial torques using the kinematics recorded from a hovering ruby-throated hummingbird [19]. This work used the kinematics in a CFD simulation, from which the wing aerodynamic forces and torques were obtained. Since the wing muscle actuation torque acted against both the aerodynamic and inertial torques, the former was calculated as sum of the latter two. There were also minor differences in the wing kinematics between the CFD study [19], and the wing and skeletal kinematic study [8], therefore, we then aligned the two kinematics and projected the wing muscle actuation torque on the wing-fixed coordinate system of the aligned wing. We then projected the actuation torques onto the body (or shoulder hinge) fixed coordinate system. These torques were finally used for optimizing the wing-actuation model parameters. Note that these actuation torques included both passive and active muscle forces.

### Optimization of wing-actuation model parameters

The unknown model parameters (including spring stiffnesses, active force and torque amplitudes, rotation slope and intercept, and muscle active periods for DMG and PiMG muscles) were optimized for each model variation to best predict the estimated wing muscle actuation torque. A full list of model parameters for each model variation, with the corresponding allowable range and the constraints for optimization, as well as relevant justifications, can be found in the Supplementary file 4 and Appendix. Genetic Algorithm (GA) (MATLAB, MathWorks, MA) was used for optimizing the parameters towards minimizing a cost function (Eq. 4) that included the following terms: i) sum of squared errors of the torques along all the three axes over the entire flapping cycle, and ii) squared error of the cumulative work at the end of the flapping cycle. The GA optimization was initialized using randomized initial values. The GA-optimized parameters were further optimized locally using interior-point optimization (implemented using fmincon function in MATLAB). We first repeated the above process four times, each time using randomized initial values. We then used the best parameter set (corresponding to the lowest cost) among the four optimized parameter sets, as the initial parameter set for the fifth optimization. The latter process was iteratively repeated 8 times to give a total of 12 runs of GA and interior-point optimization. The entire process was repeated thrice and the best parameter set among the 36 runs was used for the final results.

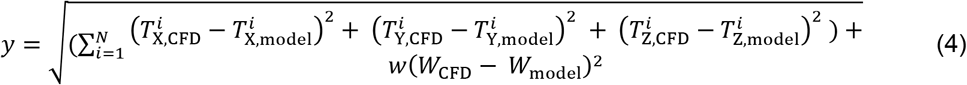

where *y* is the total cost, *i* is the time index of data points ranging from 1 to N (equal to 100) covering the complete flapping cycle, 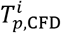 and 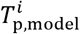 are the CFD-estimated and model-predicted torques at time index *i* about X, Y, or Z axis (Fig. A3a), and *W*_CFD_, *W*_model_ are the CFD-estimated and model-predicted cumulative work for the complete flapping cycle, respectively. The weight *w* (equal to 40) for work was empirically determined to render this term comparable to other terms in the cost function.

The optimized model parameters can be found in Supplementary file 1. Note that the final results are from the cases with the linear parallel springs for all muscles, since the model variations with nonlinear springs provided only minor improvement in prediction.

## Appendix

### Calculations of wing shoulder, elbow, and wrist angles

The shoulder, elbow, and wrist angles are defined as sequential angles of rotation with respect to the mid-downstroke orientation to match the instantaneous skeletal orientation (see Hedrick et. al. [8] for details). For calculating these angles (Fig. 1a), we followed the same method by Hedrick et. al. [8] for shoulder and elbow, and we slightly modified the method for calculating wrist angles to enable better interpretation of wrist joint functioning. Specifically, we redefined wrist axes as follows: i) W1 as the wrist chordwise axis on the humeral-ulnar plane (HUP) that is perpendicular to the wrist long-axis, ii) W3 as the wrist long-axis lying along the manus, and iii) W2 as the wrist normal axis perpendicular to W1 and W3, thereby building an orthogonal wrist coordinate system (Fig. 1a). We then estimated the rotation angles about W1 and W2 at a given flapping instant similar to those described by Hedrick et. al. [8]. Finally, the rotation angle about W3 was estimated as the angle required to align the secondary feather location for the hitherto rotated and the actual recorded wings. Note that W3 does not always lie on HUP due to wing bending and as a result, the wrist normal axis W2 may not always be perpendicular to HUP.

### Literature on muscle anatomy and functions of hummingbird wings

The studies on muscle anatomy and functions of hummingbird wings are relatively scarce compared with other avian species, however, it is known that hummingbird wings share muscle types and insertion regions similar to other avian wings [9]. The most comprehensive works to date are from Zusi et. al and Welch et. al. [9,10], where insertion points of most hummingbird flight muscles and part of their functions (a summary of which can be found in Fig. A1 and Supplementary file 3) are described. Hummingbird wing musculoskeletal system comprises of primary muscles, which are pectoralis and supracoracoideus, and secondary muscles, which are the smaller wing muscles that connect the bones of the axial skeleton (Coracoid, Clavicle, Sternum, and Scapula) to the wing bones and the wing bones among themselves [9,10]. Together, these muscles provide elaborate and rich actuation of the shoulder (3-DoF), elbow (1-DoF), and wrist (3-DoF) joints [8].

**Fig. A1.**
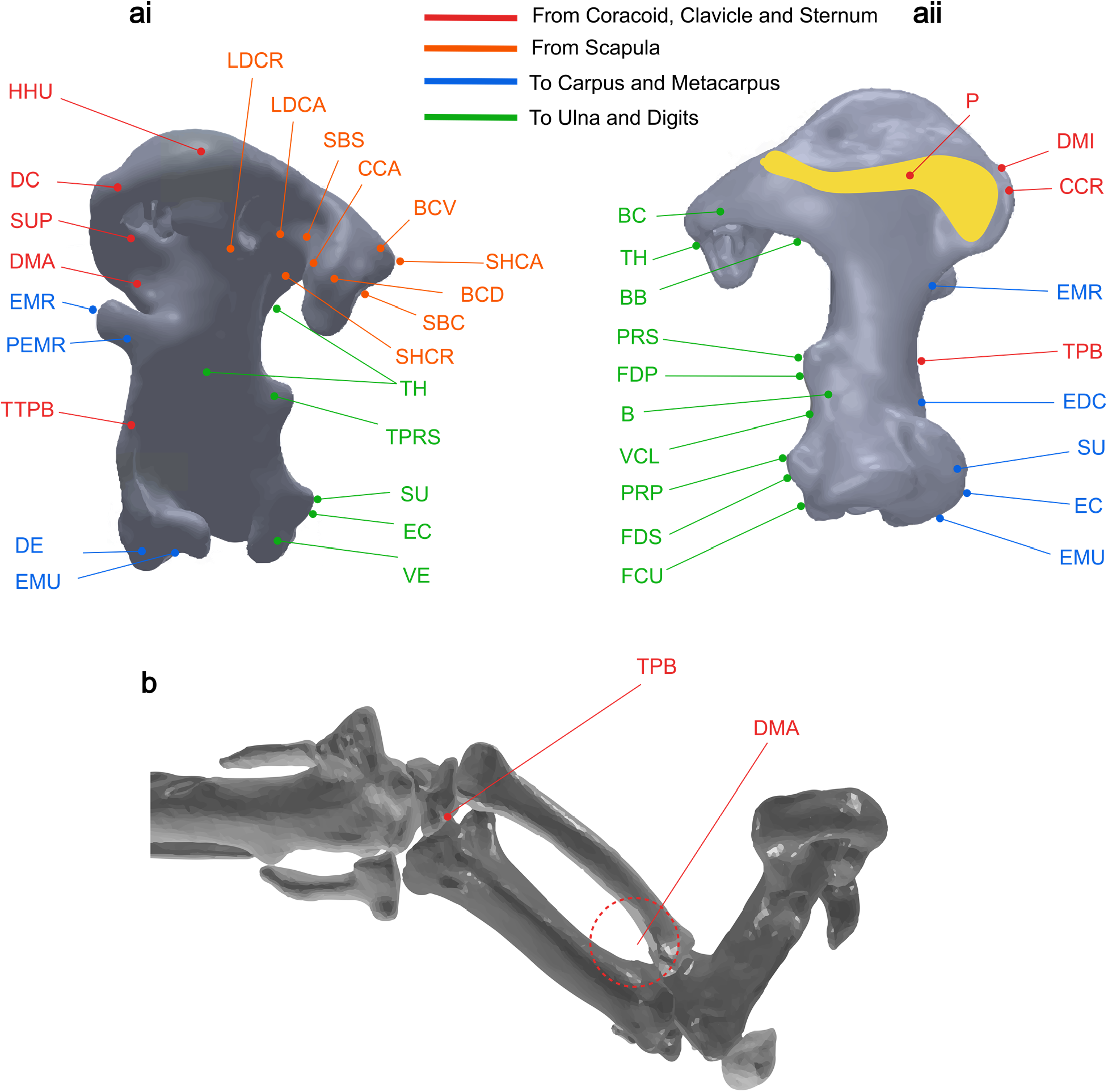
Muscle attachment points on humerus, radius, and ulna. **(ai)** Dorsal view of the humerus. **(aii)** Ventral view of the humerus. **(b)** Side view of the radius and ulna, note that only muscles originating from the body skeleton are shown. The connection points and muscle abbreviations are adopted from Zusi et. al. [9]. The images of wing skeleton bones were generated using CAD files provided by Hedrick et. al. [8].

The primary muscles are the two largest muscles that insert on the humerus. It is known that hummingbirds have evolved to leverage long-axis rotation of the humerus (the proximal wing bone that connects to the shoulder joint), which enables fast, large-amplitude stroking and power with lesser strains (proportional change in length) of the primary muscles than in other birds [8,11]. Using the available wing skeletal kinematics data of a hovering hummingbird [8], and the knowledge of insertion points of these muscles (obtained through literature and dissections, see below), we modeled the kinematics and dynamics of these muscles to determine their individual functions.

For the secondary muscles inserting on the humerus, since we could not precisely determine their individual function, we hypothesized that these muscles can collectively deviate and pitch the wing skeleton with respect to different axes (DMG_h_/PiMG_h_,DMG_w_/PiMG_w_, and DMG_b_/PiMG_b_, see Fig. 1c). These axes were defined to comprehensively cover all reasonable possibilities. We assumed that stroking effect due to these muscles was insignificant compared to that of pectoralis and supracoracoideus, since pectoralis and supracoracoideus are substantially more powerful compared to these muscles (in terms of size, pectoralis and supracoracoideus are three times larger than the third largest muscle M. scapulohumeralis caudalis [9], which in turn is larger than other muscles).

A similar anatomical study for the radius and ulna [9] revealed that the majority of the muscles originating from the body skeleton and inserting on the radius/ulna exert forces in (and parallel to) HUP (Fig. A1b). We therefore hypothesized that these muscles primarily deviate the wing skeleton. Note that the moment arms for these muscles are much larger compared to any other muscle attaching to the humerus, and therefore the torques due to these muscles can be similar or larger than those due to primary muscles, despite their relatively low forces. Together with the deviation torque created by muscles inserting on the humerus, we hypothesized a Deviation Muscle Group (DMG). Similarly, to model the pitching effect due to the muscles inserting on the humerus, we hypothesized a Pitching Muscle Group (PMG).

### PECT and SUPRA insertion point coordinates on humerus in the canonical (mid-downstroke) position

We defined the coordinates of PECT and SUPRA insertion points on the humerus in the canonical position, using the knowledge of these insertion points obtained from Zusi et. al. [9] and dissections (see below), from which all subsequent skeletal rotations, in accordance with the skeletal data from Hedrick et. al. [8], were performed. This was done by first defining these coordinates for humerus, which was oriented such that its long-axis lied along negative Y-axis (transverse body axis), with origin defined at the shoulder joint, using the dissections and the CAD model for the humerus available through Hedrick et. al. [8]. We then rotated the humerus (and the corresponding insertion points) to align its long-axis with that in the mid-downstroke position, followed by rotating the humerus about its long-axis to qualitatively fit the skeletal orientation in the mid-downstroke position as found by Hedrick et. al. [8]. The final rotated humerus (and the corresponding insertion point coordinates) position was defined as the canonical mid-downstroke position.

### Dissection experiments for insertion points of pectoralis, supracoracoideus, and the M. scapulohumeralis caudalis

We dissected the pectoral girdle of a male calliope hummingbird (*Selasphorus calliope*) to find the insertion regions for pectoralis, supracoracoideus, and the M. scapulohumeralis caudalis. We used a dissecting scope and fine forceps to reveal the insertions of the muscles on the humerus. We interpreted *in vivo* action by holding the external wing in an extended posture and pulling on a given muscle belly. We found that the pectoralis connects to the entire insertion region on the humerus as shown in yellow in Figs. A1 and A2, instead of a single insertion point as previously described by Zusi et. al. [9]. Accordingly, in the initial wing-actuation model, while we assumed that at each time instant PECT inserts on the humerus at a single effective insertion point, this point can vary linearly along the anatomical insertion region (PH_1_ to PH_2_) throughout the half-stroke, which was done by including additional parameters *p_start_* and *p_end_*. The calculation of instantaneous insertion point for PECT is represented by Eqs. 5–7,

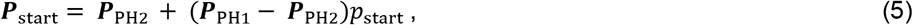

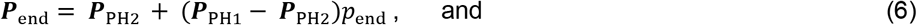

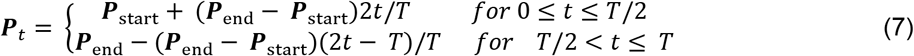

where ***P***_PH1_ and ***P***_PH2_ are the coordinates of the endpoints PH_1_ and PH_2_ on PECT insertion region, with shoulder joint as the coordinate system origin (Fig. A2), ***P**_t_*, ***P***_start_, and ***P***_end_ are the coordinates for PECT insertion point at time *t* (where *t* = 0 corresponds to the beginning of upstroke), the start of upstroke, and the end of upstroke, *T* is the flapping period, and *p*_start_ and *p*_end_ are the parameters for optimization, which can vary from 0 to 1, where 0 represents ***P***_PH2_ and 1 represents ***P***_PH1_, respectively.

We found that including the provision for variable insertion point for PECT did not improve the cost for 7 model variations (compared to when we assumed that PECT inserts at the bottommost point PH_2_), improved the cost by a maximum of only 4% for 4 model variations, and by 9.7% for one model variation. Moreover, for 3 out of 5 model variations with improved cost, both *p*_start_ and *p*_end_ were a maximum of 0.023, indicating that effectively PECT attaches at the inferior most point (PH_2_) on the humerus. Secondly, an accurate representation of the force vector could not be obtained for the instance when PECT attaches at an intermediate point between PH_1_ and PH_2_. Therefore, we assumed that PECT effectively attaches at PH_2_. Anatomically, this is justified since in the case when PECT effectively applies force at an intermediate point on the humerus, the tendon must wrap around the humerus to exit at PH_2_ and consequently the tendon is effectively attached at PH_2_ on the humerus.

**Fig. A2.**
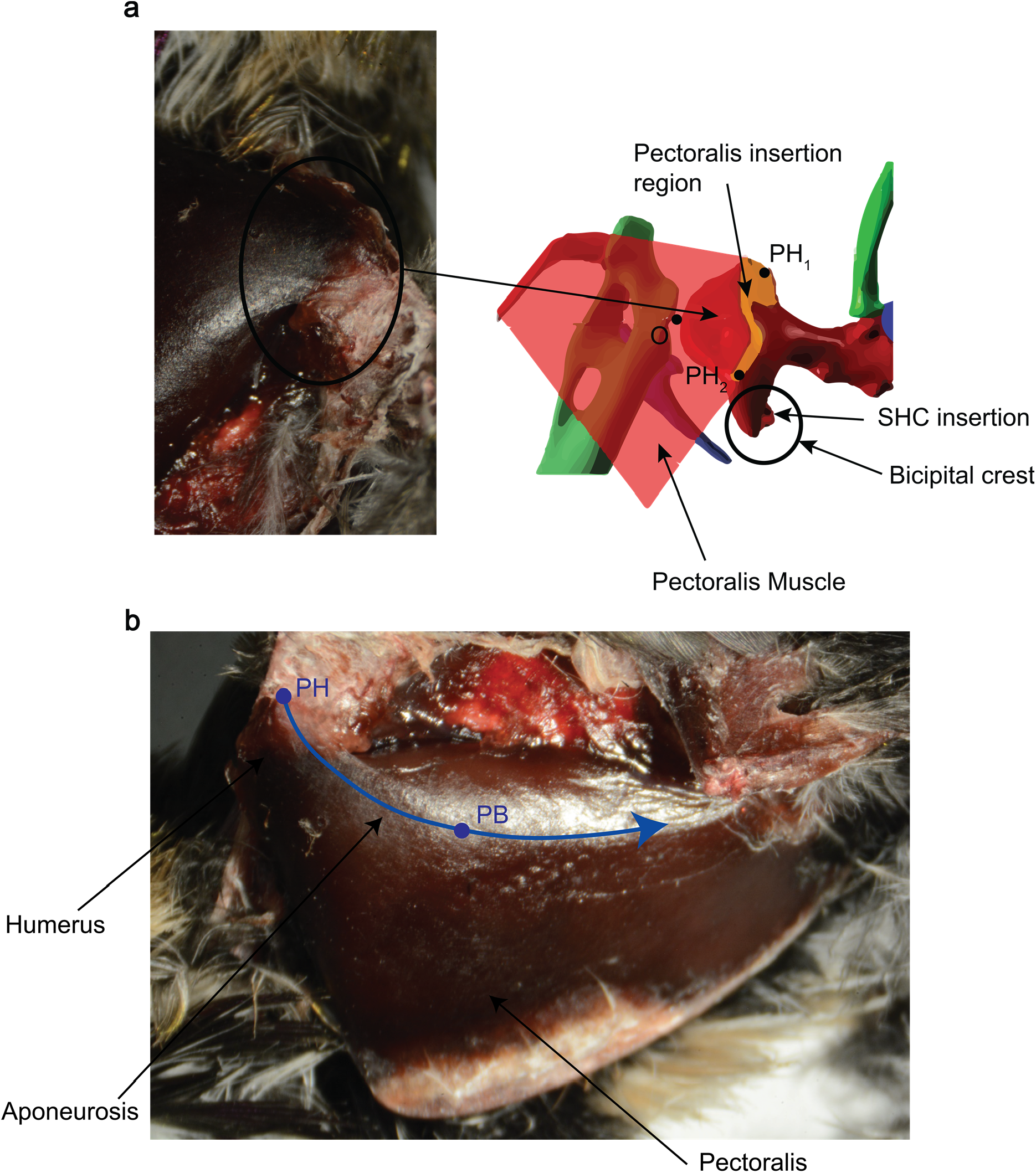
Superficial view of the left pectoralis and its insertion on the humerus in a calliope hummingbird (*Selasphorus calliope*). **(a)** Region of connection of muscle on the humerus as found through dissection, where the two extremes are marked PH_1_ and PH_2_. The coordinates of points PH_1_ and PH_2_ are defined with respect to the shoulder joint (O) as the origin. **(b)** Left view of the pectoralis. The humerus is oriented in the mid-downstroke orientation. The blue line represents the central tendon of the pectoralis. PH represents the pectoralis insertion point on the humerus and PB represents the effective insertion point of the pectoralis on the body skeleton, lying along the Aponeurosis.

### Force directions of PECT and SUPRA

The direction of PECT force can be represented by the tangent to the tendon at its humerus insertion point. We assumed that PECT force direction can vary as flapping cycle progresses, since pectoralis bends around the humerus (see Fig. A2, where the tendon (blue line) is highly bent), and the degree of bending varies linearly with time. We included additional variables to represent this changing direction of the force vector: *θ*_s_, *θ*_pz_, and *θ*_py_, which represent the instantaneous rotation angles for SUPRA in the tendon plane and for PECT about Z-axis and Y-axis as described below.

To find the direction of PECT force vector at any instant, we began by defining the nominal force direction represented by the unit vector connecting PECT insertion points on the humerus and the body skeleton, i.e., vector 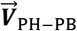. The nominal force vector can then be represented by 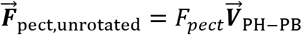 (Fig. A3a), where *F_pect_* is the force magnitude. We rotated 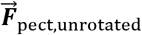 by the instantaneous rotation angle −*θ*_pz_ (details follow in the next paragraph) about Z-axis to get 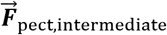, followed by a rotation about Y-axis by *θ*_py_ to give the final rotated PECT force vector 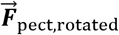. We similarly rotated SUPRA force vector from its initial orientation 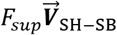 (Fig. A3b), by *θ_s_* in the plane of SUPRA tendon.

The instantaneous rotation angles *θ*_py_, *θ*_pz_, and *θ*_s_ are linear functions of flapping time and are represented by Eq. 8.

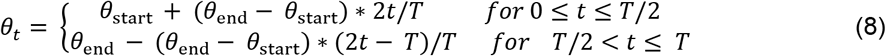

where *θ_t_* is the instantaneous rotation angle (either of *θ*_s_, *θ*_pz_, and *θ*_py_) at time *t* (where *t* = 0 corresponds to the beginning of upstroke), *θ*_start_ and *θ*_end_ are the rotation angles at the beginning and the end of the upstroke, and *T* is the flapping period, respectively. The parameters *θ*_start_ and *θ*_end_ were set as parameters for optimization. The flapping period *T* was set to 23.2 milliseconds as found by Hedrick et. al. [8] for ruby-throated hummingbirds.

The instantaneous length of PECT or SUPRA was estimated using the corrected direction of muscle force vector. The muscle was assumed to lie along a circular arc that lies along the muscle’s aponeurosis and connects the effective insertion points on the body skeleton and the humerus, as shown in Figure A3d. The length of the arc was calculated using Eq. 9:

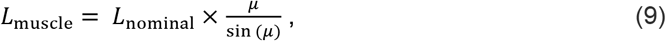

where *L*_nominal_ and *L*_muscle_ are the instantaneous distance between insertion points of the muscle and the calculated muscle length, and *μ* is the angle between nominal and rotated force vectors, respectively.

**Fig. A3.**
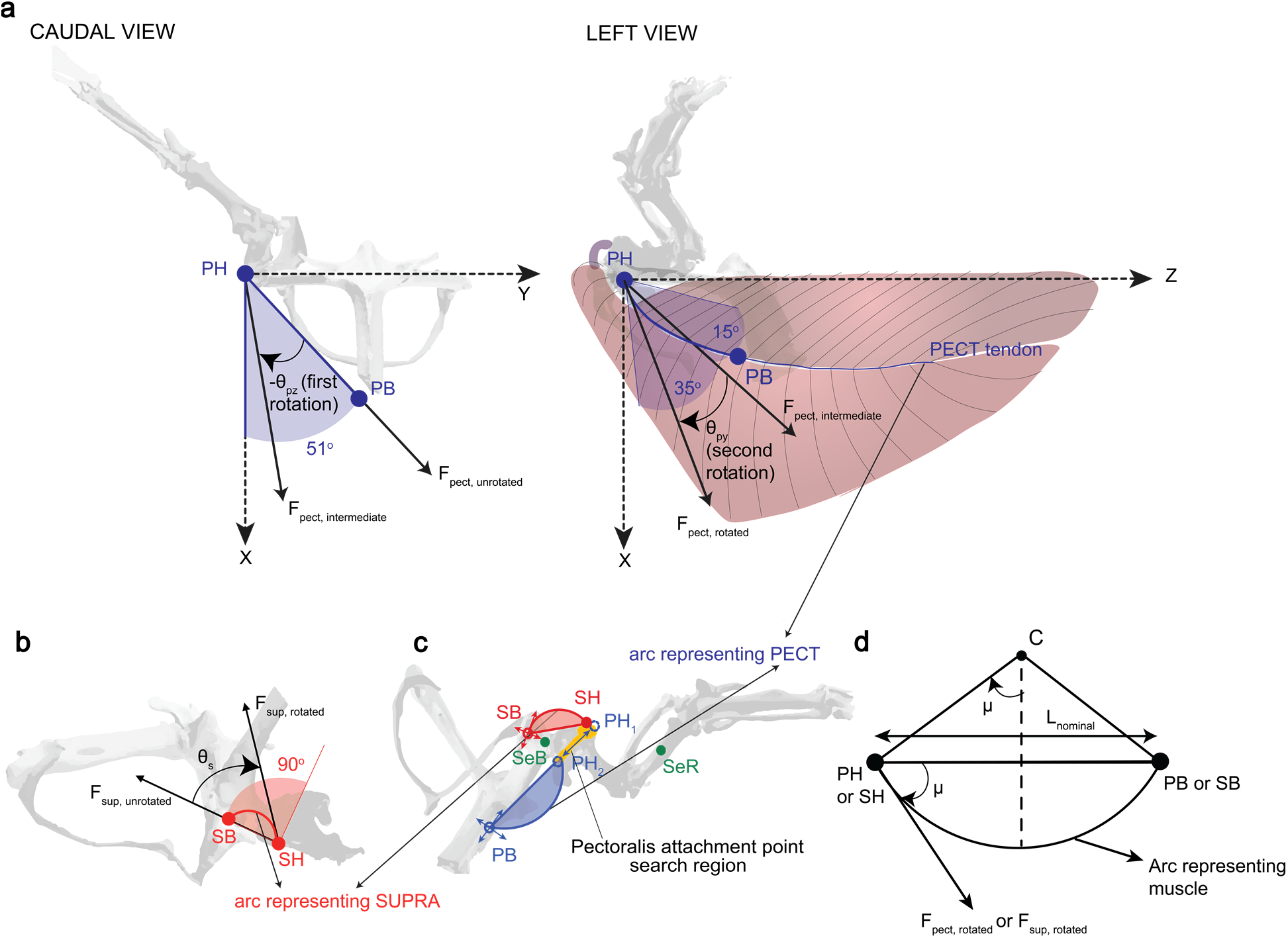
Summary of the method for finding the instantaneous direction of PECT and SUPRA force vector, PECT and SUPRA muscle length, muscle insertion points, and L_WL_. **(a)** PECT: First, the nominal force vector ***F***_pect,unrotated_ (along straight-line PH-PB) was rotated about Z-axis by −θ_pz_ to give ***F***_pect,intermediate_, followed by a second rotation about Y-axis by θ_py_ to give ***F***_pect,rotated_. The maximum possible rotation about Z-axis is 51°, and the minimum and maximum possible rotation about Y-axis are - 15° and 35° (see blue shaded regions for range). **(b)** SUPRA: It was assumed that SUPRA tendon lies instantaneously in a plane defined perpendicular to the Y-Z plane and containing the straight-line SH-SB. The rotated force vector ***F***_sup,rotated_ was found by rotating the nominal force vector ***F***_sup,unrotated_ (along straight-line SH-SB) by an angle θ_s_ in the plane of the tendon. The maximum possible rotation is 90° (see red shaded region for range). **(c)** Effective insertion points or regions of PECT and SUPRA on body and humerus, and the effective insertion points of combined secondary muscles (used for estimating L_WL_). Each muscle insertion point is named after the muscle (group) name (i.e., P for PECT, S for SUPRA, and Se for secondary muscles), followed by the bone that the muscle inserts on (i.e., H for humerus, R for radius, and B for body skeleton). A filled circle represents a fixed insertion point with known location, an open circle (with double-sided arrows) represents a fixed insertion point with unknown location (on body skeleton, being searched in optimization), and a dotted open circle represents an effectively time-varying insertion point along a defined insertion region. PH_1_ and PH_2_ represent the two extremes of PECT insertion point on the humerus. An example of PECT (blue) and SUPRA force (red) vectors and the corresponding muscle arcs for length calculation are shown. (**d**) Pictorial description of the arc length calculation, where the arc represents the muscle that lies along the aponeurosis, ***L***_nominal_ is the straight-line distance between insertion points of the muscle and ***μ*** is the angle between nominal and rotated force vectors, respectively (see Eq. 9 for details).

### Modeling active muscle force (or torque) profile using a central pattern generator

To model the active force (or torque) generation (i.e. contractile component) in the muscle models (Fig. 1c), we aimed at identifying a mathematical model that: 1) replicates an actual muscle’s contractile force profile and 2) can be modulated such that the output can be shaped by external commands (such as signal from brain or sensors) to enable simulation of different aspects of flapping flight. We chose the Matsuoka central pattern generators because:

i. They have well-defined mechanisms for including external commands to modulate amplitude, phase, and frequency of its output [75].
ii. They can generate a rich set of output profile waveforms to capture the possible muscle force (or torque) profiles [75]. It can be seen that the CPG output profiles, in general, mimic an actual muscle’s contractile component force profile more closely than simple harmonic functions.
iii. They have been successfully used to generate and/or control gaits in biologically inspired locomotion such as snakes [76], salamanders [77], bipedal [78], and quadrupedal locomotion [79].

#### Active force (or torque) profile

First, we generated CPG output for a given constant input *s_i_* using the set of differential Eqs. 10–14 representing the Matsuoka two-neuron CPG [80]. The coefficients in these equations were empirically determined for generating sustained oscillations at 43.08 Hz, the flapping frequency of Ruby-throated hummingbird [8], and can be found in Table A1. Feedback can also be incorporated in the equations based on the multiple neuron CPG model by Nakamura et. al. [81].

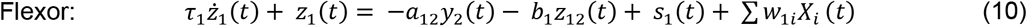

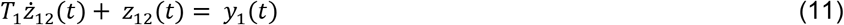

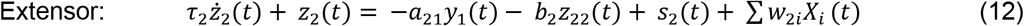

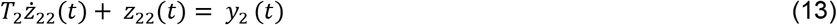

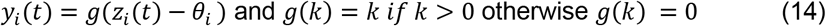

where *t* is the time, *z_i_* and *y_i_* are the state and output of the *i^th^* neuron, *z*_*i*2_ is the internal inhibition state for *i^th^* neuron, *τ_i_* and *T_i_* are the time constants for *z_i_* and *z*_*i*2_, *α_ij_* (i ≠ j) is the inhibition coefficient for the complementary output, *b_i_* is the inhibition coefficient for *z*_*i*2_, *w_ji_* are the feedback gains for *j^th^* neuron and sensory feedback *X_i_*, *θ_i_* is the offset for output and is set to zero, and *s_i_* is the input to the *i^th^* neuron, respectively.

**Table A1:**
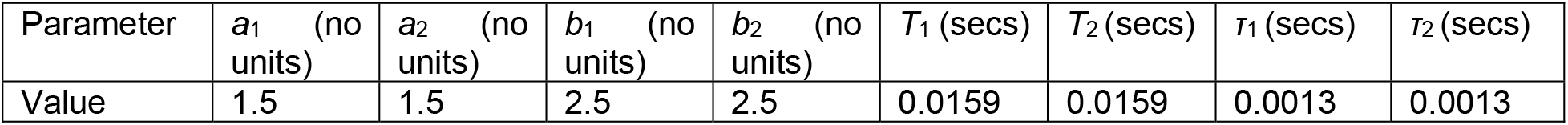
CPG parameters for generating periodic output at 25 Hz.

#### Active force (or torque) onset timing and period

First, we describe the active force (or torque) period for a muscle, i.e., the time interval for which the muscle active force (or torque) is non-zero. The pectoralis force generation in hummingbirds is known to span an entire half-stroke [82]. However, active force periods for other muscles are not known. The supracoracoideus force in pigeons is known to span almost an entire half-stroke [32] and in this work, we assumed that SUPRA active force spans the entire half stroke for a ruby-throated hummingbird. However, for each of the combined secondary muscles, we included an additional parameter to represent the active muscle torque period. The parameter lied between zero and one, where zero represents that active torque starts and ends at the same time and one represents that active torque spans the entire half-stroke.

Next, we describe the active force (or torque) onset timing. The onset timing of active forces by pectoralis, supracoracoideus, and some of the secondary muscles in the hummingbird wing musculoskeletal system are reported by Altshuler et. al. [49], however, no data is available for other muscles. It was found that the onset of supracoracoideus active force happens at about half of the flapping cycle period after the onset of pectoralis force [49]. Accordingly, we fixed the time period between PECT and SUPRA active force onsets. To account for the possibility that the onset of active force due to PECT or SUPRA and the onset of the respective half stroke do not overlap, an additional parameter *Φ* (active force onset phase) was included for optimization (see Supplementary file 4 for more details).

The final active force profile for a given muscle for the given active force period and onset timings was estimated by first shifting the active force (spanning the entire half cycle) using Eq. 15:

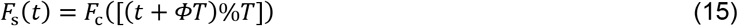

where *F*_c_ and *F*_s_ are the original and shifted force profiles, *t* represents the time variable, *Φ* is the PMG onset phase, % represents the modulo function, and *T* is the flapping cycle period, respectively. This is followed by changing the time scale to reduce the muscle active force period while keeping the amplitude same. Note that for PECT and SUPRA, since the active force periods span the entire half-stroke, no such change in time scale was required.

### Final wing muscle actuation torque estimation

The torque 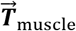 due to PECT or SUPRA for a given insertion point 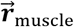 on the humerus (with origin at the shoulder joint) and given force vector 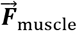 was found using the cross product,

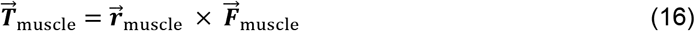

### Modeling M. scapulohumeralis caudalis (SHCA)

M. scapulohumeralis caudalis is the third largest muscle in hummingbirds and hence may contribute significantly towards wing actuation in all the three degrees of freedom. The insertion point for M. scapulohumeralis caudalis was found through the dissection and additional parameters for direction correction for SHCA, similar to PECT, were included for optimization. We found that including SHCA did not improve model prediction since the optimized SHCA torque direction coincided with that of DMG and PiMG, resulting in redundancy. Therefore, SHCA was not included explicitly in the final model variations.

### Estimation of torque due to DMG and PiMG

DMG and PiMG were modelled to represent the effect of muscles other than PECT and SUPRA as described in the previous sections. To estimate the torque vector due to each of these muscle groups, we first estimated the group’s instantaneous torque direction i.e. *DMG_i_* and *PiMG_i_*, where *i* represents wrist, humerus, or body (Fig. 1c and 1d), using instantaneous skeleton coordinates. We then estimated the group’s torque magnitude using the three-component model, whose output represents torque rather than force. We modelled torque directly rather than force since the moment arms of any of the individual muscles, which these muscle groups represent, were not known. Similar to PMG, we assumed that the onset times of active torques due to the two muscles in each of the secondary muscle groups were spaced out by a half-stroke period. Note that the active torques due to the two muscles may not be antagonistic, i.e. they might be applied in the same direction, which is not known beforehand. We therefore included two binary variables during optimization representing the directions of the active torques due to each of the two muscles. In summary, a group’s torque magnitude was estimated using Eq. 17:

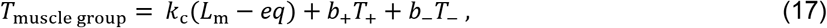

where *T*_muscle group_ represents the torque magnitude due to a muscle group (DMG or PiMG), *k_c_* represents the total stiffness of the muscle group, *L*_m_ represents the muscle length measure (see Fig. 1d), *eq* represents the measure of equilibrium position of the muscle group, *T*_+_ and *T*_−_ represents the active torques due to antagonistic muscles, and *b_+_* and *b*_−_ represents the binary variables representing the directions of active torques, respectively. The active torque profiles were found using Eq. 15, similar to the active force profiles of PMG, using the optimized parameters for DMG or PiMG (amplitude, muscle active periods, and active torque onset phase, i.e., *ϕ*_DMG_ or *ϕ*_PiMG_ when it was independent of that of PMG and *Φ* when it was identical to that of PMG). The non-linear torque models were represented using the equations similar to Eqs. 1–3.

### Estimation of wing muscle actuation torque using CFD data

The wing muscle actuation torque can be defined as the total torque transferred from bird’s body to its wing, applied at the shoulder. It overcomes both the wing inertia and the aerodynamic pressure to move the wing, as described by Song et. al. [19].

We first estimated the wing muscle actuation torque trajectory for a single wing flapping cycle (Fig. 1b) based on the computational fluid dynamics (CFD) study by Song et. al. [19] (referred to as ‘CFD study’), where CFD simulations were performed using the wing kinematics recorded from an actual hovering hummingbird and from which both the wing aerodynamic and inertial torque were obtained. We then projected it to the wing and skeletal kinematics data obtained by Hedrick et. al. [8] (referred to as ‘wing and skeletal kinematics study’), where skeletal kinematics along with wing kinematics were measured using micro-CT and X-ray methods. The kinematics used in both these studies were recorded in the ‘wing and skeletal kinematics study’ on different ruby-throated hummingbirds (*Archilochus colubris*). Since the studies were conducted using data from different birds, there exist minor differences in wing kinematics in the two studies. We synthesized the two sources of data by assuming that the muscle actuation torque relative to the wing frame were identical, thereby enabling us to project the muscle torque data from the ‘CFD study’ to the ‘wing and skeletal kinematics study’ in the wing frame. The following describes the detailed synthesis procedure of the two data sources:

#### i) Estimation of wing muscle actuation torque using CFD data

Song et. al. [19] conducted CFD simulations for a hovering ruby-throated hummingbird based on kinematics recorded in the ‘wing and skeletal kinematics study’. They divided the wing area into 615 mesh elements and estimated aerodynamic forces (and torques) on each of these elements as the flapping progressed. In addition, they also estimated the averaged moment of inertia of the wing (averaged over a flapping cycle), which was used to estimate the inertial torque of the wing at each instant. Based on the obtained aerodynamic and inertial torques, they estimated the total wing muscle actuation torque that was transferred from the bird’s body to its wing using Eq. 18.

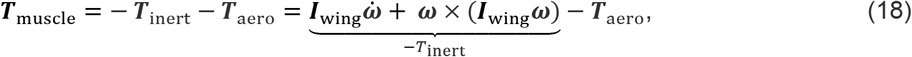

where ***I***_wing_ is the averaged wing moment of inertia, 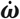 and ***ω*** are the wing angular acceleration and the angular velocity, ***T***_muscle_ is the instantaneous wing muscle actuation torque, −***T***_inert_ is the torque required to overcome instantaneous wing inertia, and ***T***_aero_ is the instantaneous aerodynamic torque experienced by the wing, respectively. All quantities are expressed in the wing-fixed frame. The CFD simulations were conducted for 7 full flapping cycles. Accordingly, we estimated ***T***_muscle_ for the 7 flapping cycles and the results were averaged to obtain ***T***_muscle_ for a single flapping cycle.

#### ii) Time alignment of wing kinematics between ‘CFD study’ and ‘wing and skeletal kinematics study’

As the first step to synthesize data from the two sources, it was required that the wing kinematics data are aligned in time, i.e., different instants of the flapping cycles in the wing kinematics from the two sources should overlap. This was achieved by first averaging the wing kinematics recorded for the 7 flapping cycles for each of the two studies, to obtain the averaged wing kinematics for a single flapping cycle. The wing kinematics from each source was then resampled at 100 times the flapping frequency. Finally, the time instants corresponding to the beginning of upstroke were matched for both the data sources to obtain the time-aligned wing kinematics.

#### iii) Spatial alignment of wing position

After the temporal alignment of wing kinematics, we observed that the resultant wing kinematics do not overlap spatially. While the aerodynamic torque, inertial, and consequently the wing muscle actuation torque were calculated based on the wing kinematics in the ‘CFD study’, it was required that these torques were projected on the wing kinematics corresponding to the ‘wing and skeletal kinematics study’, for more accurate estimation of the wing muscle actuation torque. At each instant of flapping cycle, we therefore shifted the wing position from the ‘CFD study’ such that it matches as closely as possible with the wing position in the ‘wing and skeletal kinematics study’. For achieving this, we first identically defined the wing-fixed planes for wings in both studies. We then shifted the wing position obtained from the ‘CFD study’ such that the wing pitching axes match, followed by a rotation about the matched pitching axis such that the wing chords corresponding to the secondary feather match as closely as possible, to give us the spatially aligned wings.

We also calculated Euler angles for the shifted HUP where the Euler angles were defined similar to that in the study by Song et. al. [83]. These Euler angles were used to calculate the rotation matrix from the wing-fixed frame to the body-fixed frame, as described in (v) below.

#### iv) Projection of CFD-estimated torques to the shifted wing

Since the two data sources correspond to the same species of hummingbird, we expected only minor differences in the overall torque characteristics. We therefore assumed identical muscle torque in the wing-fixed frame, and the wing-fixed inertial, aerodynamic, and wing muscle actuation torques, as obtained from the ‘CFD study’, were projected on the wing-fixed axes of the shifted wing.

#### v) Estimation of wing torque in the body-fixed frame

Given the instantaneous torque vector in the wing-fixed frame (***T***), the torque vector in the body-fixed frame (***T***_b_) was obtained using Eqs. 19 and 20,

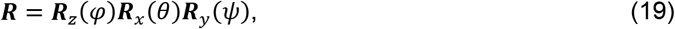

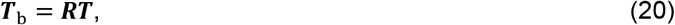

where ***R*** is the rotation matrix from the wing-fixed frame to the body-fixed frame, ***R**_z_, **R**_x_*, and ***R**_y_* are the rotation matrices for stroke, deviation, and pitching axes and *φ, θ*, and *ψ* are the stroke, deviation, and pitching angles, respectively.

#### vi) Wing muscle actuation torque represented as a series of Euler torques

The Euler torques were calculated using the Eqs. 21 and 22. These equations were derived by extracting the components of body-fixed torque ***T**_b_* along the stroke, deviation, and pitching axes.

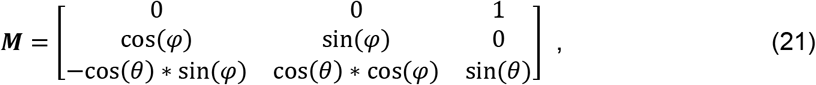

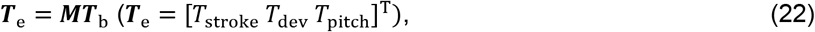

where ***T***_e_ is the Euler torque vector, *T*_stroke_, *T*_dev_, and *T*_pitch_ are the stroking, deviation, and pitching torques, and ***M*** is the matrix used for extracting ***T***_e_ from ***T***_b_, respectively. Note that the Euler angles are different for the wing-fixed frame of reference and the HUP frame of reference. Accordingly, the Euler torques are different for the wing and wing skeleton.

### Cumulative work calculation

The cumulative work done by PECT or SUPRA muscle force was calculated by integrating the dot product of the displacement and the muscle force over time. The total work done by the wing muscle actuation torque was obtained similarly using the data from the ‘CFD study’. For muscles other than PECT and SUPRA, since individual muscles were not modelled separately and only lumped equivalent length was used for the two muscles in DMG or PiMG, we were unable to directly calculate the cumulative work done by individual muscles of the secondary muscle groups. Instead, we estimated the cumulative work done by DMG and PiMG combined using the difference between the total cumulative work (done by the total wing muscle actuation torque) and the work done by PECT and SUPRA.

### Percentage error calculation between actual and model torque

We first calculated the absolute error between the actual and model-predicted torque (for each of the 12 model variations) at each instant of flapping cycle (i.e., at 100 time instants spanning a complete flapping cycle). We then took the mean of the absolute errors at these 100 time instants for each of the stroking, deviation, and pitching torques, and the percentage error was calculated with respect to the maximum absolute actual stroking, deviation, or pitching torques, respectively.

### Constraints enforced while parameter optimization and justification

Supplementary file 4 summarizes the different parameters included in the optimization for all twelve model variations, their lower and upper limits, and corresponding justifications. The following describes additional constraints between some of the parameters that were enforced to ensure realistic parameter values:

1. Parallel spring stiffness is always smaller than series spring stiffness for both PECT and SUPRA. This is because the tendon is assumed to be stiffer compared to muscle [67].
2. 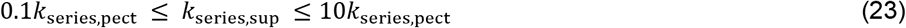

where *k*_series,pect_ is the series spring stiffness for PECT and *k*_series,sup_ is the series spring stiffness for SUPRA. This is to ensure reasonable values for the spring stiffnesses.
3. 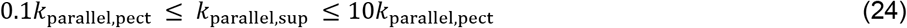

where *k*_parallel,pect_ is the parallel spring stiffness for PECT and *k*_parallel,sup_ is the parallel spring stiffness for SUPRA. This is to ensure reasonable values for the spring stiffnesses.
4. *θ*_s,max_ ≥ *θ*_s,min_ This is because we assumed that SUPRA is in relaxed state at the end of upstroke and consequently the length of SUPRA is higher at the start of upstroke then at the end of upstroke, which is proportional to the instantaneous angle of rotation.
5. *θ*_pz,max_ ≥ *θ*_pz,min_ and *θ*_py,max_ ≥ *θ*_py,min_ This is because physically it can be seen that due to the orientation of the bicipital crest of the humerus, pectoralis’s force vector orients more ventrally at the beginning of downstroke compared to the end of downstroke.

### Calculation of stress and mass-specific power for the primary muscles

For calculating the muscle stress (kPa) at any instant of flapping cycle, we divided the muscle force by *Acosα*, where *A* is the muscle area and *α* is the pennation angle as found through dissection of a calliope hummingbird (17° for pectoralis and 30° for supracoracoideus). To calculate *A*, we assumed pectoralis (0.29 g) and supracoracoideus (0.12 g) mass as found through dissections conducted earlier as a part of the Hedrick et. al. study [8] for a ruby-throated hummingbird. We then calculated muscle volumes (pectoralis = 0.27 cm^3^; supracoracoideus = 0.12 cm^3^) using the standard muscle density 1060 Kg m^−3^ [86]. We then calculated the average fascicle lengths (pectoralis = 8.5 mm, supracoracoideus = 4.85 mm) using the scaled figures provided in Welch et. al. [10]. Finally, the averaged cross-sectional area was calculated by dividing the muscle volume by the respective average fascicle length and multiplying by cos*α*. This provided the estimate for the physiological cross-section area of the muscles as 0.30 cm^2^ for pectoralis and 0.21 cm^2^ for supracoracoideus.

The mass-specific power for PECT and SUPRA was calculated by dividing the instantaneous muscle power by the respective muscle mass.

## Acknowledgments

This research was supported by the Office of Naval Research (ONR) (Program Officer: Dr. Marc Steinberg, award number: N00014-19-1-2540 to B.C., B.W.T, and H.L.). We would also like to thank Dr. Jialei Song for providing guidance on CFD data interpretation.

## Data and code availability

All code and data for preprocessing, model optimization, plotting, and analysis are available on GitHub repository (https://github.com/ska5623/Hummer_musculoskeletal_modelling) and Dryad (doi:10.5061/dryad.n5tb2rbxg).

## Description of Supplementary files

**Figure 5 – figure supplement 1:** Active and passive torque components from PECT, SUPRA, and combined secondary muscle groups for stroke **(a)**, deviation **(b)**, and pitching **(c)** axes (as defined based on HUP, Fig. 1a) for **(i)** body-centered and **(ii)** elbow-centered models. Two torque scales are used: N-mm and the product of body weight (Mg) and wing length (R). The solid curves show the median torque components (median torque at each time instant estimated using the torque results for the four body-centered or elbow-centered model variations) and the associated colored shaded areas represent the region spanned by torques for the four body-centered or elbow-centered model variations. The gray shaded and unshaded areas represent downstroke and upstroke, respectively.

**Supplementary file 1**: **Optimized parameters for all the wing-actuation model variations.** The vertical axis range represents the range set during optimization, unless specified otherwise. **(a)** Relative amplitude of the CPG outputs, which is proportional to the amplitude of PECT and SUPRA active forces. **(b)** Equilibrium position of the two primary muscles, where 0 represents the minimum muscle length during the flapping cycle. **(c)** Stiffness of parallel elastic components of PECT and SUPRA. **(d)** Stiffness of series elastic components of PECT and SUPRA. **(e)** Relative stiffness of the DMG and PiMG muscle groups. Since we used relative measures for muscle length for these muscle groups, the exact muscle stiffness could not be found. **(f**) Active time period for DMG and PiMG muscles, which represents the % half-stroke for which the active torque is >0 for a given muscle. **(g)** Relative amplitude of the four CPG outputs (for two muscles of DMG and two muscles of PiMG), which are proportional to the amplitude of their respective active torques. Note that the two muscle active torques in each of DMG and PiMG could have acted along the same direction. **(h)** Time instant at which DMG and PiMG were at their respective equilibrium positions (i.e. when the muscle group’s passive force is zero), represented by the flapping cycle instant, where 0% represents the beginning of upstroke. **(i)** Relative muscle group active force onset phase for each of the PMG, DMG, and PiMG. Note that six variations had aligned active force onset phases for all muscle groups, while for the other six variations, the active force onset phases were different for different muscle groups. **(j)** Effective insertion points of PECT and SUPRA on the body skeleton. Note that these are not the actual muscle endpoints. **(k)** Starting and ending angles for the different rotation angles, used for correcting PECT and SUPRA force directions. Note that the possible range during optimization are different for different angles and are shown on the right. **(l)** Percent error in stroking, devition, and pitching torques calculated against the maximum torque (see details in the text).

**Supplementary file 2**: Different model predicted muscle functioning variables for each of the 12 model variations (Fig. 1d).

**Supplementary file 3:** Description of insertion points and function of various hummingbird muscles. **Supplementary file 4:** Description of parameters that were optimized for each of the 12 musculoskeletal model variations, their lower and upper limits, and justification for these limits.

**Movie S1:** Left view (from the left towards the right wing) of the hummingbird, demonstrating the instantaneous wing skeleton movement, the force vectors due to PECT (blue) and SUPRA (red), and arcs representing muscles (along with the straight line connecting the effective muscle insertion points on body skeleton and wing skeleton). The lengths of force vectors are proportional to their respective magnitudes. The humerus is shown in red, the radius and ulna are shown in blue, and the manus is shown in orange. A total of two flapping cycles are shown where the second cycle repeats the first one and illustrates the key hummingbird wing musculoskeletal actuation principles (Fig. 6). The illustration is shown at the beginning and the middle of the half-strokes, where the red, blue, and green straight or curved arrows in the intermittent illustrations represent the torque due to SUPRA, PECT, and combined secondary muscles, and the lengths of the arrows are proportional to torque magnitudes; the black curved dotted arrows represent the direction of wing pitching movement. The three figures on the left shows the instantaneous stroke, deviation, and pitching torques (marked by black dots) due to active and passive elements of PECT, SUPRA, and the combined secondary muscle groups; the solid curves show the median torque components (median torque at each time instant estimated using the torque results for the four wrist-centered model variations) and the associated colored shaded areas represent the region spanned by torques for the four wrist centered model variations.

**Movie S2:** Cranial view (viewed from bird’s head towards tail) of the skeletal movement and primary muscles forces and kinematics, along with the wing actuation torque in stroke, deviation, and pitching directions, shown only for a single flapping cycle. Bird’s eyes are not shown for clarity.

**Movie S3:** Dorsal view of the skeletal movement and primary muscles forces and kinematics, along with the wing actuation torque in stroke, deviation, and pitching directions, shown only for a single flapping cycle.

